# Pathogenic T-bet^+^ memory B cells worsen multiple sclerosis through myeloid cell activation

**DOI:** 10.64898/2026.09.01.748682

**Authors:** Rajiv W Jain, Maria Goiko, Antonio Mendes, Sara Xue, Salma Zein, Rianne Gorter, Marlene T Mørch, Alexandre Prat, Rui Li, V. Wee Yong

## Abstract

Depletion of B cells in multiple sclerosis (MS) is beneficial yet there is no defined pathogenic B cell subset in MS. Here, we demonstrate that T-bet^+^ memory B cells are rare in control human brain specimens but are found in MS lesions. Transfer of murine T-bet^+^ memory B cells into mice with ongoing experimental autoimmune encephalomyelitis (EAE), a model of MS, worsened their disability and demyelination. Microglia/macrophage density increased in central nervous system parenchyma even though B cells mostly remained in barriers. In culture, secreted factors from T-bet^+^ memory B cells promoted macrophage migration. IFN-γ produced by T-bet^+^ memory B cells activated microglia to secrete chemokines that further enabled macrophage migration. Consistent with their age-associated elevation, conditional deletion of T-bet in B cells lowered EAE severity in aged but not young mice. We define T-bet^+^ memory B cells as pathogenic in EAE and MS through their IFN-γ-facilitated interactions with microglia/macrophages.

## Introduction

A major cause of neural injury in multiple sclerosis (MS) is neuroinflammation driven by microglia and infiltrated immune cells such as monocyte-derived macrophages^1^. Indeed, MS lesions are often classified by the organization of microglia and macrophages where those with confluent coverage are labelled active lesions, those with microglia/macrophages concentrated at the edge as mixed active/inactive (or chronic active), and those without immune cells as inactive^2^. Inflammation is also commonly found within the meninges of post-mortem MS brains frequently associated with adjacent grey matter injury and accumulation of microglia/macrophages^3,4^. There is considerable evidence of the detriments of microglia/macrophages in MS through their production of toxic molecules^5^, induction of demyelination^2,6^, and the activation of T cells^7^. Factors responsible for switching normally-protective microglia to a pathogenic state are of much interest^5^.

B lymphocytes are also crucial in MS as their depletion using monoclonal antibodies targeting CD20 is one of the most effective therapies for treating relapsing disease^8,9^. While B cells are found in MS lesions^10^, their abundance is variable between lesions and patients^11,12^. B cells are rare in the parenchyma but can accumulate in large numbers in CNS borders such as perivascular spaces or meninges^4,13,14^. In the meninges, B cells can form organized B cell follicle-like structures that are correlated with the accumulation of macrophages in the meninges and microglia in the underlying parenchyma and with increased expression of inflammatory proteins such as iNOS or TNF-α^4^; these changes may occur through proteins secreted from B cells^10,15^.

What has complicated interpretations of B cell pathophysiology is that there are many different subsets of B cells^10,16^. Developing B cells are detected in the cerebrospinal fluid (CSF) of MS patients^17^, possibly derived from skull bone marrow^18,19^; there are also elevated numbers of B1 and B2 lineage mature naïve B cells^20^. Memory B cells, referring to those that participated in an immune response that then became quiescent^21^, or B cells that acquire effector functions^22^, are also present in larger numbers in the CSF of MS patients^23–25^. Germinal center B cells that undergo somatic hypermutation to yield high-affinity B cell progeny^26^ have been detected in the CNS of animal models of MS^27^; their presence in MS lesions has not been confirmed and is controversial^10,11,20^. Terminally differentiated B cells, known as antibody-secreting plasma cells or plasmablasts, are concentrated in the CSF^20^ and lesions^28^ of MS. Regulatory B cells are also detected in the CNS of people with MS^29,30^.

Effective MS therapies directed at B cells affect the memory B cell pool, while ineffective drugs or those that worsen disease do not^31,32^. Memory B cells are highly diverse; two major types are conventional and age-associated B cells (ABCs), the latter also known as atypical memory B cells. Conventional memory B cell subsets have traditionally been defined based on their differentiation fates during secondary immune responses^21^ with some poised to rapidly form germinal centers or differentiate into plasma cells^33^. These cells accumulate in the CSF in MS patients^23^, particularly during active disease^24^. Atypical memory B cells, discovered more recently^34,35^, have been linked to anti-pathogen immune responses^36,37^, aging^34^, and autoimmune disorders^38,39^. They are typically T-bet^+40^ and are commonly identified by the surface markers CD27^-^ CD21^-^, CD11b^+^, or CD11c^+^ B cells^41^. Despite evidence using human samples suggesting that memory cells are pathogenic in MS, no B cell or memory B cell subset has been proven to be pathogenic in animal models. Here, we provide evidence for the pathogenicity of T-bet^+^ memory B cells in the EAE animal model and MS, and we describe their production of IFN-γ that coordinates the activation and recruitment of microglia and macrophages to mediate neuropathology.

## Results

### T-bet^+^ memory B cells are found in MS lesions

Although previous studies have shown T-bet^+^ memory B cells are increased in the CSF and brains of MS patients^42,43^, whether these cells accumulate within MS lesions is unknown. We first sought to address this using postmortem brain samples from individuals with MS. We stained brain tissues from 5 non-neurological disease controls and 7 MS cases (Supplemental table 1) for CD20^+^ T-bet^+^ cells (Figure 1). In control meninges or parenchymal tissue, CD20^+^ B cells were rare and were not detected in the white matter. A single CD20^+^ cell was found in the meninges of 3 of 5 control tissues where one of these was T-bet^+^ (Figure 1B).

**Figure 1.**
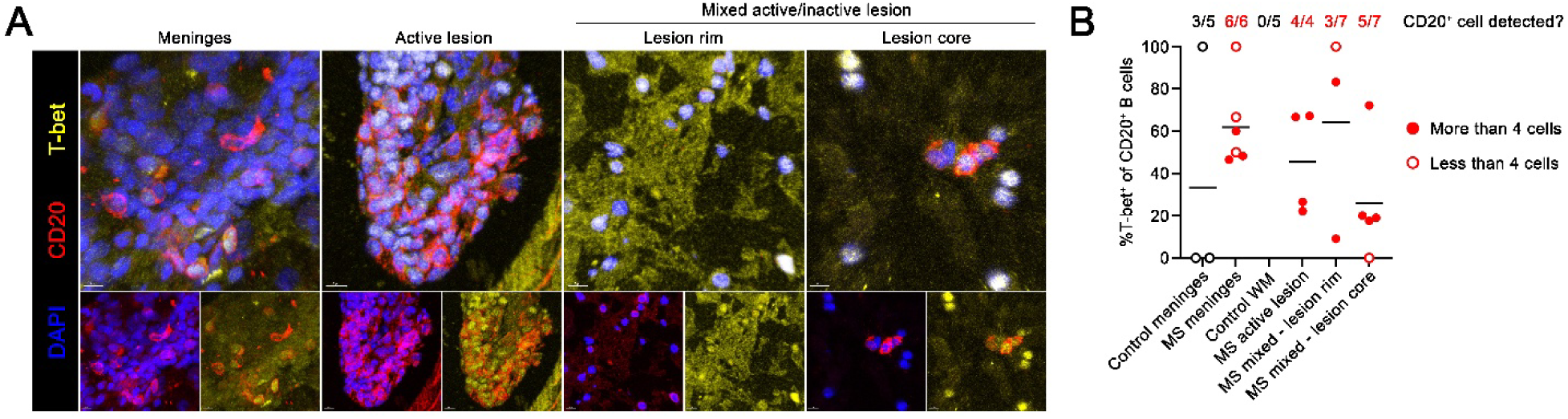
T-bet^+^ memory B cells are found in MS lesions. (A) Representative images depicting T-bet expression in CD20^+^ B cells in various MS lesion types. Scale bars are 8 μm. (B) Quantification of the percentage of CD20^+^ B cells that are T-bet^+^ across the indicated lesion types. The proportion of tissues analyzed that had at least 1 CD20^+^ B cell is indicated at the top and measurements based on 1-3 B cells are indicated through hollow circles; full circles represent measurements with greater than 4 B cells (number of lesions analyzed per type: n = 4-7). See also Figure S1.

Next, we assessed MS brain sections and classified lesion types based on the absence of PLP staining and the pattern of HLA-DR staining (Supplemental figure 1A). CD20^+^ cells were much more frequent than in control tissues, and were found in the inflamed meninges of 6 of 6 MS cases, and in 4 of 4 active lesions, the rim of 3 of 7 mixed active/inactive lesions, and the core of 5 of 7 mixed active/inactive lesions (Figure 1A and 1B). Of these, nearly every single lesion had a T-bet^+^ CD20^+^ B cell except for one; however, the last had only 2 CD20^+^ cells detected and thus could be from under sampling. Most of the B cells imaged were found in perivascular cuffs (Supplemental figure 1B), although some rare T-bet^+^ B cells were found in the parenchyma. Overall, T-bet expression in border-associated (meningeal and perivascular) B cells is common across MS lesion types although the presence of B cells within each specimen is variable.

### Transferred T-bet^+^ memory B cells worsen disability in mice with ongoing B cell-independent EAE

We developed an *in vitro* polarization protocol to generate large numbers of B cells with defined properties. Naïve B cells extracted from young mice were maintained in a naïve state or polarized into classical memory B cells or T-bet^+^ memory B cells with no differences in viability (Supplemental figure 2A). Polarization into T-bet^+^ memory B cells resulted in 98.2% of cells expressing T-bet relative to 1.69% and 1.39% seen in naïve or classical memory B cell cultures, respectively, using a Tbx21-tdTomato reporter mouse^44^ (Figure 2A). Of significance, while CD95 expression was very low on naïve B cells (7.86%), 74.6% and 97.6% were expressed on polarized classical memory B cells and T-bet^+^ memory B cells, respectively, indicative of their activation.

**Figure 2.**
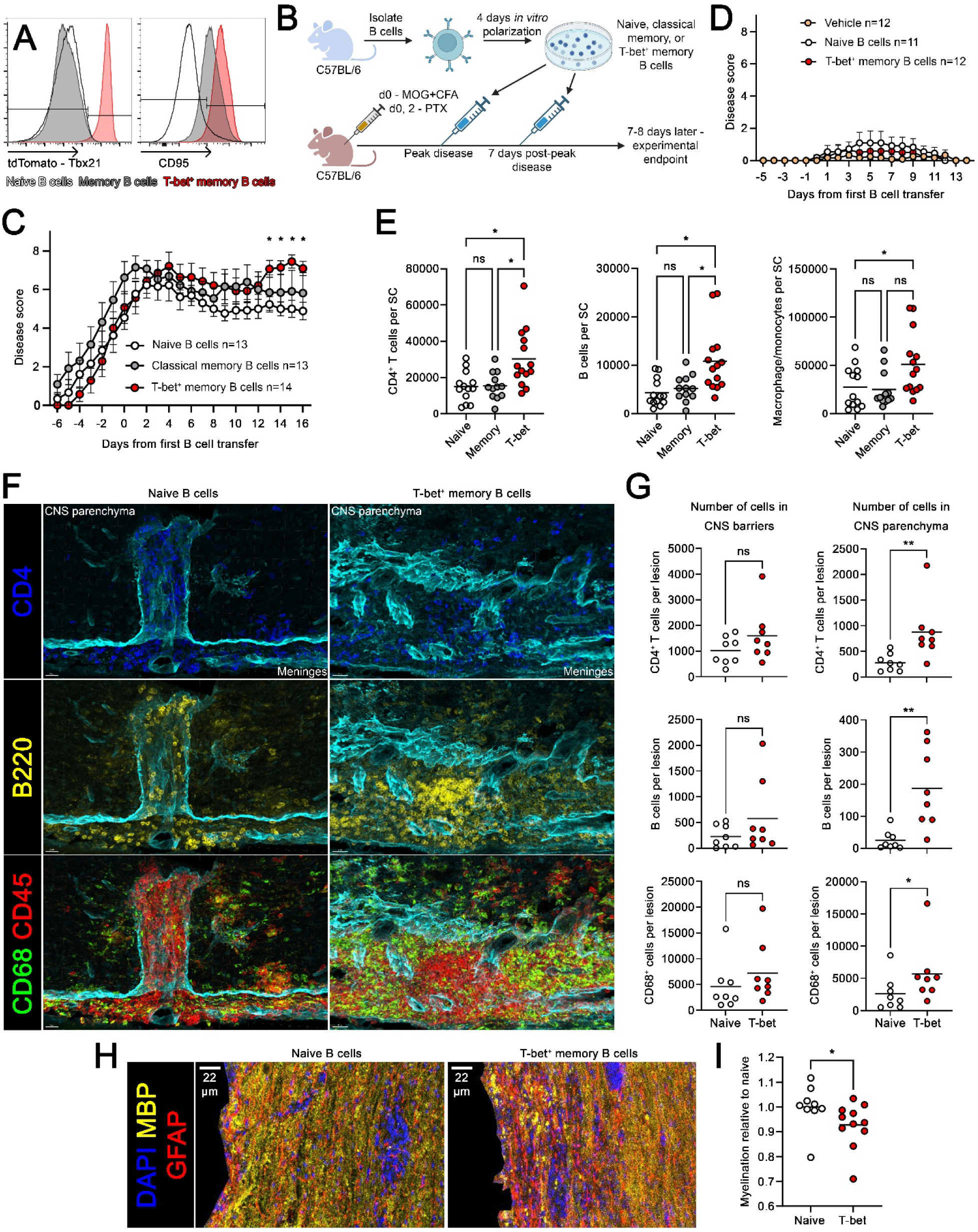
Transferred T-bet^+^ memory B cells worsen disability in mice with ongoing B cell-independent EAE. (A) Histograms showing the expression of Tbx21 using a transcriptional reporter tied to tdTomato expression and CD95 expression amongst *in vitro* polarized B cells detected by flow cytometry. (B) Schematic showing the process used to test B cell pathogenicity in MOG_35-55_ EAE mice, and the timing of intravenous transfer of *in vitro* polarized B cells at EAE peak disease and seven days later. (C) EAE disease score curve of the experiment described in B. Combination of 3 independent experiments (n = 13-14). (D) EAE disease score curve of an experiment similar to that in B except without pertussis injection. Combination of 2 independent experiments (n = 11-12). (E) Quantification of the absolute numbers of CD4^+^ T cells, B cells, and Ly-6C^+^ macrophages in the spinal cord of the experiment shown in C (n = 12-14). (F and G) Representative images of histology from the spinal cord of mice with transferred naïve or T-bet^+^ memory B cells, with the number of CD4^+^ T cells, B cells, or CD68^+^ macrophage/microglia in CNS barriers or in the CNS parenchyma quantified on a per lesion basis. Representative 1 of 2 experiments (n = 8). Scale bars are 20 μm. (H and I) Representative images showing DAPI, GFAP and MBP staining to demonstrate demyelination in spinal cord, where demyelination standardized to naïve B cell transfer is quantified. Combination of 2 independent experiments (n = 9-11). Scale bars are 22 μm. Statistical significance was assessed using a two-way ANOVA with Geisser-Greenhouse correction and Tukey’s post-hoc test (C and D), a one-way ANOVA using Kruskal-Wallis test and Dunn’s post-hoc test (E), a Mann-Whitney U test (G and I). Data are presented as mean ± SEM (C and D) or mean (E, G, and I) and p-values are summarized as *p<0.05 and **p<0.01. See also Figure S2.

To test whether T-bet^+^ memory B cells are pathogenic *in vivo*, we used MOG_35-55_ induced EAE in 8-12 weeks young mice as B cells are not pathogenic in this model^45–47^, so that the activity of transferred B cells could be assessed. At the peak of EAE disability and seven days after, mice were intravenously infused with *in vitro* polarized B cell subsets (Figure 2B). Remarkably, the transfer of T-bet^+^ memory B cells worsened the already severe disability (a score of 3 on the traditional 5-point scale), and this was not observed for the transfer of naïve or classical memory B cells (Figure 2C).

In another experiment where MOG_35-55_ EAE and adjuvant was administered but without pertussis toxin injections to cause suboptimal EAE, the transfer of T-bet^+^ memory B cells did not induce disease (Figure 2D). These results suggest that T-bet^+^ memory B cells amplified pathogenicity when EAE is established rather than initiating the disease process (Figure 2D).

Analyses of plasma by multiplex Luminex showed that mice with transferred T-bet^+^ memory B cells had elevation of numerous inflammatory factors relative to naïve B cell transfer or unimmunized mice, including VEGFA, EPO, M-CSF, GM-CSF, IFN-β1, IFN-γ, TNF-α, IL-1β, IL-2, IL-3, IL-7, IL-9, IL-10, IL-11, IL-120p70, IL-13, IL-15, CXCL9, CXCL10, CCL3, CCL4, CCL17, CCL19, CCL21, and LIF (Supplemental figure 2B). Flow cytometry (Supplemental figure 2C) revealed that mice with transferred T-bet^+^ memory B cells had increased numbers of CD4^+^ T cells, B cells, and Ly-6C^+^ macrophages in their spinal cord (Figure 2E) but not increases of neutrophils or CD8^+^ T cells (Supplemental figure 2D).

As increases in immune cells could reflect these cells accumulating in CNS barriers without entering the CNS parenchyma, we used multiplex staining coupled with single cell analysis of EAE lesions using histoflow cytometry^48,49^ to quantify cell abundances and cellular localization (Supplemental figure 2E and 2F). The numbers of CD4^+^ T cells, B cells, and CD68^+^ myeloid cells increased within the CNS parenchyma of EAE mice with transferred T-bet^+^ memory B cells (Figure 2F and 2G) but not in laminin^+^ CNS barriers. Staining for MBP revealed that T-bet^+^ memory B cell transfer was also associated with increased demyelination (Figure 2H and 2I). Overall, T-bet^+^ memory B cells cause injury in mice with ongoing EAE, associated with increased disease burden and immune cell recruitment into the CNS parenchyma.

### Transferred T-bet^+^ memory B cells accumulate in the spleen and CNS during EAE

We determined where transferred T-bet^+^ memory B cells accumulate during EAE. Mb1-cre mice (with a CD79A promoter driven cre-recombinase^50,51^) were crossed to Ai9 mice^52^ to generate mice with tdTomato specifically expressed in B cells (Supplemental figure 3A). TdTomato^+^ B cells were then polarized into naïve or T-bet^+^ memory cells and transferred into wildtype EAE mice (Supplemental figure 3B) and subsequently analyzed by histoflow cytometry (Supplemental figure 3C) or flow cytometry (Supplemental figure 3D).

TdTomato^+^ T-bet^+^ memory B cells accumulated in the spleen and CNS of EAE mice but not within inguinal lymph nodes relative to naïve B cell transfer (Figure 3A-3C). Transferred T-bet^+^ memory B cells mostly maintained T-bet expression in all analyzed organs and few endogenous B cells (tdTomato^-^) were found to be expressing T-bet across these tissues. TdTomato^+^ T-bet^+^ memory B cells were predominately found on the outside of the spinal cord or in circular clumps (Figure 3A) suggesting they accumulated in the meninges and perivascular spaces, respectively. Nonetheless, some tdTomato^+^ T-bet^+^ memory B cells were seen in the parenchyma. Some transferred T-bet^+^ memory B cells differentiated into germinal center B cells (CD38^lo^ CD95^hi^) or plasma cells in the spleen but otherwise they maintained a phenotype consistent with memory/naïve B cells in the CNS and lymph nodes. Thus, T-bet^+^ memory B cells concentrated in the CNS and spleen during EAE and they differentiated into other B cell subsets specifically in the spleen.

**Figure 3.**
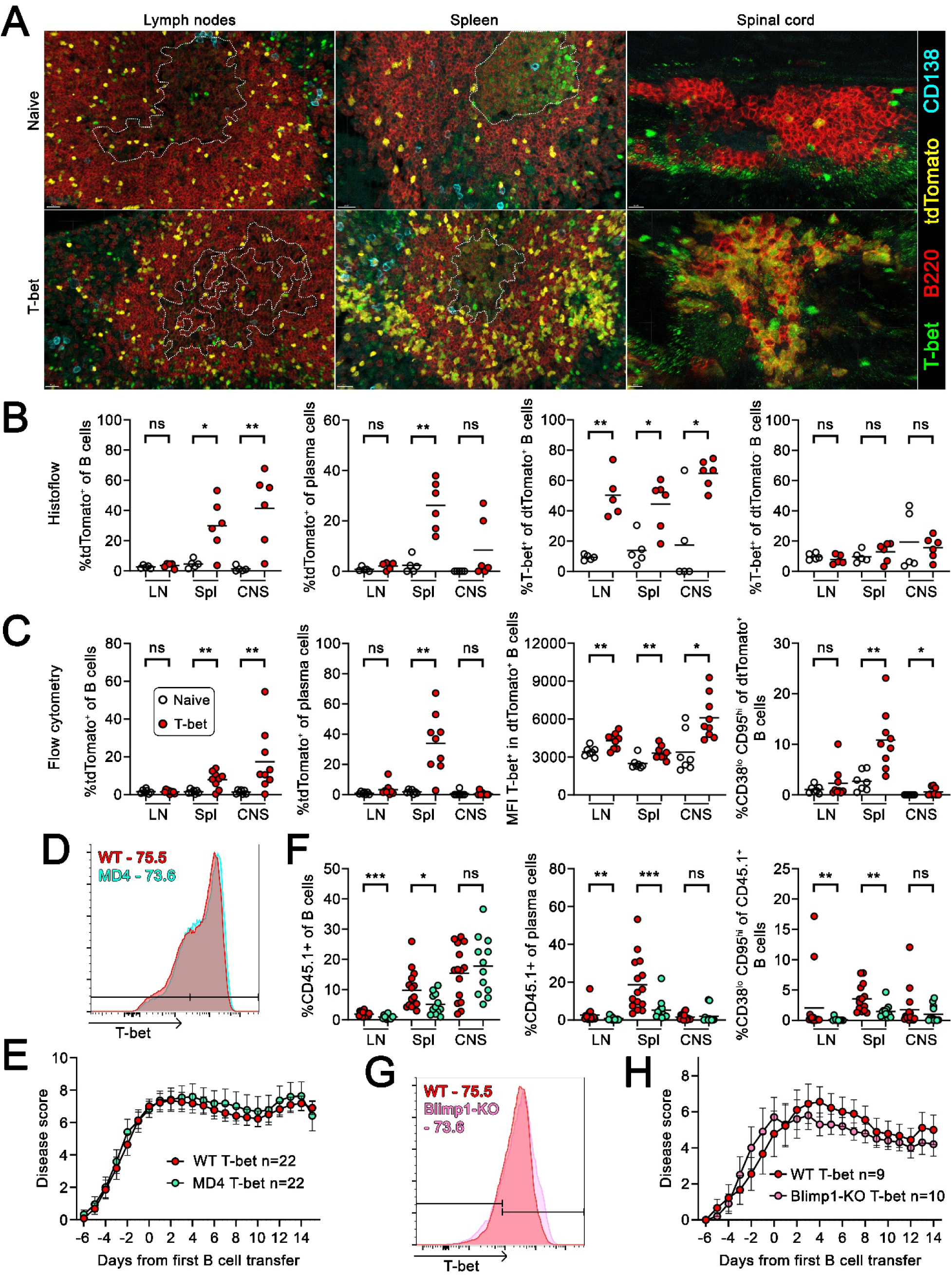
T-bet^+^ memory B cell pathogenicity is independent of autoreactivity or plasma cell differentiation but is associated with CNS accumulation. (A and B) Transfer of tdTomato^+^ (derived from Mb1^cre^ x Ai9 mice) *in vitro* polarized B cells in EAE mice was used to track B cell accumulation in the inguinal lymph nodes (LN), spleen (Spl), and spinal cord (CNS) by histology and quantified by histoflow cytometry (n = 5-6). Scale bars are 20 μm for LN and Spl, 10 μm for CNS. Dashed lines outline germinal centers based on reduced IgD staining (not shown). (C) Quantification of the identical experiment as above by flow cytometry. Representative of 1 of 2 experiments (n = 7-9). (D) Histogram showing T-bet expression in *in vitro* polarized CD19^+^ B220^+^ B cells from wildtype or MD4 mice. (E) EAE disease score curve comparing CD45.2^+^ EAE mice with transferred *in vitro* polarized T-bet^+^ memory B cells from CD45.1^+^ or CD45.1^+^ MD4 mice. Combination of 4 independent experiments, with one using CD45.2^+^ transferred B cells (n = 22). (F) Quantification of transferred B cell infiltration into the LN, Spl, and CNS and the differentiation status of the EAE mice from E using flow cytometry. Combination of 3 independent experiments (n = 12-15). (G) Histogram showing T-bet expression in *in vitro* polarized CD19^+^ B220^+^ B cells from wildtype or T-bet^cre^ x Blimp1^flox^ (Blimp1-KO) mice. (H) EAE disease score curve comparing EAE mice with transferred *in vitro* polarized T-bet^+^ memory B cells from wild type or Blimp1-KO mice. Combination of 2 independent experiments (n = 9-10). Statistical significance was assessed using single or multiple Mann-Whitney U tests (B, C, E, F, and H). Data are presented as mean ± SEM (E and H) or mean (B, C, and F) and p-values are summarized as *p<0.05, **p<0.01, and ***p<0.001. See also Figure S3.

### Autoreactivity and plasma cell differentiation are not required for T-bet^+^ memory B cell pathogenicity

As germinal center responses are driven against antigens^26^, and no foreign antigen was provided, this suggested that the transferred T-bet^+^ memory B cells may be recognizing autoantigens in the spleen thereby contributing to their pathogenicity by expanding autoreactive B cell clones or through the production of autoantibodies. To test whether autoreactivity impacted the capacity of transferred T-bet^+^ memory B cells to affect EAE, we used the MD4 mouse strain^53^ in which all B cells have fixed specificity for the hen egg lysozyme. B cells from CD45.1^+^ wild type or CD45.1^+^ MD4 mice were *in vitro* polarized into T-bet^+^ memory B cells (Figure 3D), then transferred into CD45.2^+^ EAE mice and analyzed. Transfer of MD4 or wild type T-bet^+^ memory B cells did not differ in their impact on EAE disease scores (Figure 3E) and there was no difference in the number of B cells, CD4^+^ T cells, macrophages, neutrophils, or plasma cells found in the CNS (Supplemental figure 3E). When analyzing where CD45.1^+^ transferred T-bet^+^ memory B cells could be detected, we found no difference in recruitment to the CNS; however, there were reduced numbers of these cells in the spleen and lymph nodes of mice with transferred MD4 T-bet^+^ memory B cells (Figure 3F). Similarly, there was decreased differentiation of plasma cells and germinal center B cells in the spleen and lymph nodes but not the CNS of EAE mice receiving MD4 cells. Together, these suggest that autoreactivity, and by extension the capacity of T-bet^+^ memory B cells to participate in peripheral germinal centers to form additional autoreactive plasma cells and B cell clones, was not required for their pathogenicity.

It was a possibility that plasma cells generated after transfer could be impacting EAE through antibody independent mechanisms^54^. To test this, we transferred *in vitro* polarized T-bet^+^ memory B cells derived from wild type or T-bet^cre55^ x Blimp1^flox/flox56^ mice (Figure 3G), in which T-bet^+^ memory B cells cannot form plasma cells as Blimp1 is required for plasma cell differentiation^57^, into EAE mice. Wild type and Blimp1-knock out T-bet^+^ memory B cells did not differ in their capacity to affect EAE scores (Figure 3H) and there was no difference in the number of immune cells found in the CNS (Supplemental figure 3F). Thus, plasma cell differentiation was not required for T-bet^+^ memory B cells to affect EAE severity.

### Transfer of T-bet^+^ memory B cells increases activation and number of macrophages and microglia in EAE lesions

Across our experiments we observed consistent increases in macrophages accumulating within the CNS suggesting T-bet^+^ memory B cells were modifying the innate immune response. As T-bet^+^ memory B cells could be affecting other myeloid cells, we made use of the CX3CR1^CreER-eYFP58^ x Ai9 mouse line to better distinguish microglia and macrophage^59^. Here, CX3CR1-CreER-eYFP x Ai9 mice were injected with tamoxifen for 5 days to label all CX3CR1^+^ cells (CNS and peripheral macrophage populations) with tdTomato. Then, 16 days later, when tdTomato was lost in peripheral monocytes due to their rapid regular turnover while long-lived CNS macrophage populations maintained marker expression (Figure 4A and 4B), EAE was induced in these mice with subsequent transfer of naïve or T-bet^+^ memory B cells (Supplemental figure 4A). Harvested spinal cords were stained with a combination of DAPI, Iba1 to identify microglia/macrophages, and endogenous tdTomato to distinguish myeloid populations. Histoflow cytometry revealed that the transfer of T-bet^+^ memory B cells, relative to naïve B cell transfer, increased accumulation of Iba1^+^ tdTomato^+^ microglia (and other long-lived CNS macrophage populations) and Iba1^+^ tdTomato^-^monocyte-derived macrophages in EAE lesions (Figure 4C and 4D, Supplemental figure 4B). To investigate their properties, we conducted unbiased clustering of single macrophage and microglia based on the expression of CD68, MHC2 and dectin-1 as activation markers; arginase-1 (Arg1) as a regulatory indicator, and Ki67 for proliferation (Supplemental 4C and 4D). Microglia and macrophages outside of lesions consisted primarily of populations (cluster 2 for microglia and cluster 14 for macrophages) with low expression for assayed markers suggesting they were mostly quiescent (Supplemental figure 4E). There were no changes in non-lesional macrophage proportions and a decrease in population 4 microglia around lesions induced by T-bet^+^ memory B cells. Within lesions promoted by T-bet^+^ memory B cells, there was a shift of cluster 2 microglia (low expression of all markers) to cluster 9 microglia with high expression of all markers (Figure 4E). Similarly, there was a shift from cluster 14 macrophages with low expression of all markers to cluster 3 macrophages that had higher level of all markers except Ki67, suggesting a general shift towards activated but non-proliferating macrophages.

**Figure 4.**
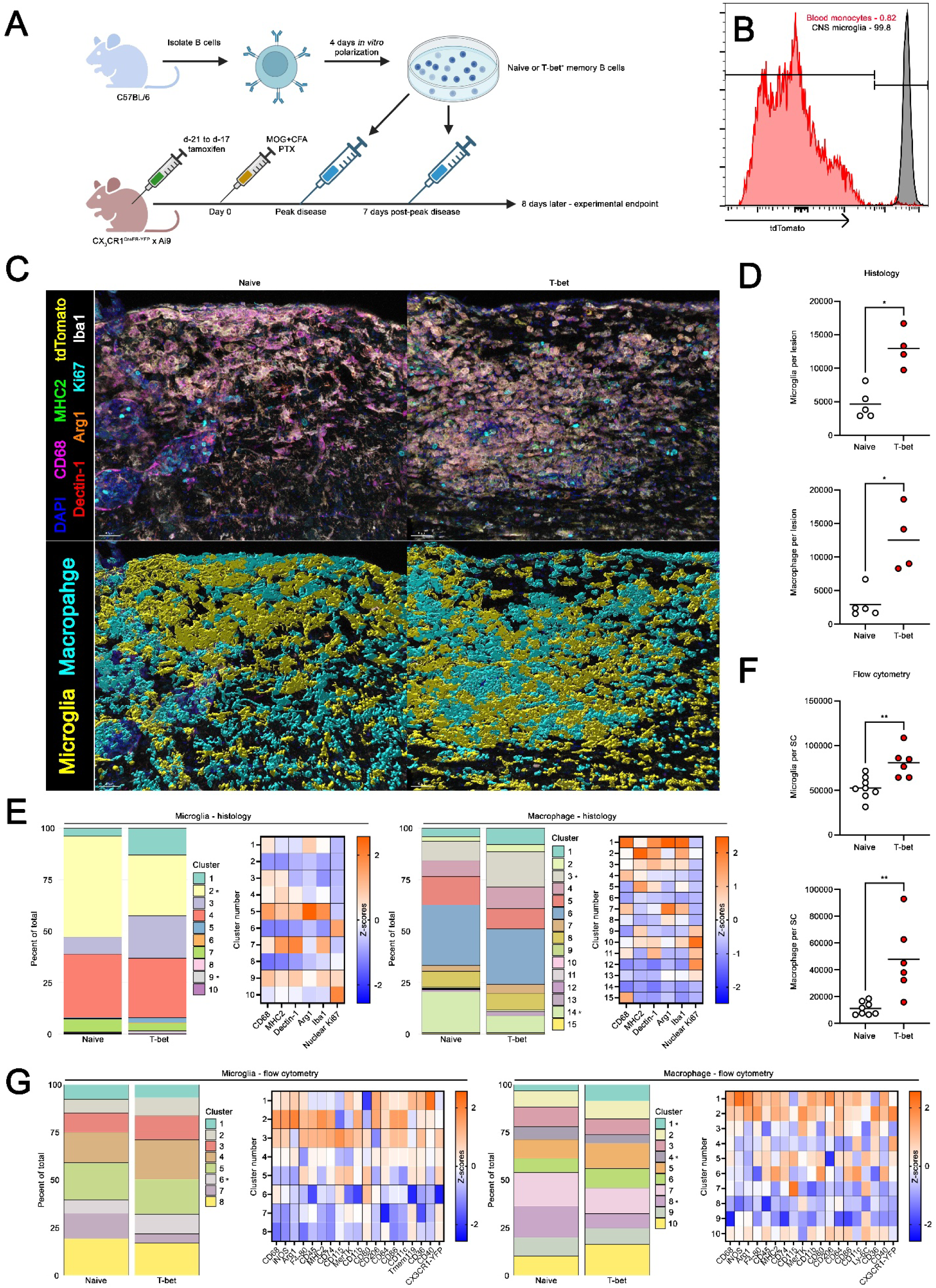
Transfer of T-bet^+^ memory B cells is associated with increased activation and numbers of macrophage and microglia in EAE lesions. (A-G) CX_3_CR1^CreER-YFP^ x Ai9 mice were injected with tamoxifen for 5 days then 16 days later, when tdTomato was retained in microglia but not peripheral macrophages (B), EAE was induced. Naïve or T-bet^+^ memory B cells from wild type mice were then injected at peak disease and 7 days later, and the mice sacrificed 8 days after the last B cell transfer to analyze their spinal cords by histoflow cytometry (C-E) or flow cytometry (F and G). (C) Representative fluorescent images (top) and mapped macrophage and microglia (bottom) around EAE lesions in the spinal cords of mice receiving naïve or T-bet^+^ memory B cells are shown and the number of each population per lesion are quantified in D (n = 4-5). Scale bars are 40 μm. (E) The proportions of microglia (left) and macrophage (right) populations by histoflow cytometry are shown along with a Z-scored heatmaps demonstrating the properties of the distinct populations. (F) Quantification of the absolute number of microglia and macrophage in the spinal cords of EAE mice receiving naïve or T-bet^+^ memory B cells using flow cytometry (n = 6-8). (G) The proportions of microglia (left) and macrophage (right) populations analyzed by flow cytometry are shown along with Z-scored heatmaps demonstrating the properties of the distinct populations. Representative one of two experiments. Statistical significance was assessed using Mann-Whitney U tests (D, E, F, and G). Data are presented as mean (D and F) or are only shown as proportion (E and G) and p-values are summarized as *p<0.05 and **p<0.01. See also Figure S4.

To corroborate these results, we used a spectral flow cytometry panel to assess microglia/macrophage properties in greater detail. We used a gating strategy to eliminate CD3^+^ T cells, NK1.1^+^ NK cells, B220^+^ CD19^+^ B cells, Ly6G^+^ neutrophils and Lyve1^+^ border-associated macrophage^60^ to obtain pure populations of CD11b^+^ tdTomato^+^ microglia and CD11b^+^ tdTomato^-^monocyte-derived macrophages for analysis (Supplemental figure 4F). Quantification of the absolute numbers of these cells reveals that both microglia and macrophages were increased in the spinal cords of mice injected with T-bet^+^ memory B cells (Figure 4F) consistent with our histology. Unsupervised clustering based on CD68, iNOS, Arg1, F4-80, CD45, MHC2, CD74, CD115, MerTK, CD11b, CD80, CD206, CD64, CD86, CD11c, Tmem119/Ly-6C, CD36, CD40, and CX3CR1-YFP revealed 8 microglia and 10 macrophage clusters (Supplemental Figure 4G). The proportion of microglia clusters was similar between EAE mice receiving naïve or T-bet^+^ memory B cells, with a small increase in the proportion of microglia with higher expression of CD68, CD80 and CD115 along with lower expression of Tmem119; these results suggest some degree of activation amongst this population (Figure 4G). Transfer of T-bet^+^ memory B cells was associated with a shift away from macrophage clusters 4 and 8 characterized by relatively low expression of most markers to cluster 1 defined by high expression of nearly all markers; these suggest a shift towards activation that combines inflammatory and regulatory markers. Similar analysis conducted on monocytes from the blood of these mice (Supplemental figure 4H) revealed no changes in the proportion of blood monocyte populations between the groups (Supplemental figure 4I and 4J) suggesting T-bet^+^ memory B cells impacted macrophage populations locally in the CNS.

### Secreted factors from T-bet^+^ memory B cells orchestrate microglia and macrophage activation and macrophage migration

We used a tissue culture paradigm to address whether T-bet^+^ memory B cells could control the observed macrophage accumulation in the CNS parenchyma and the local activation of microglia/macrophages. B cells were polarized *in vitro* into naïve, classical memory, or T-bet^+^ memory B cells for 3 days, then totally replenished with new culture media containing only BAFF as a supplement. Their respective supernatants were collected after 24 hours (Figure 5A). Supernatants or media containing BAFF were then placed below transwell filters and bone marrow-derived macrophages (BMDMs) were added on top of the filters; the number of BMDMs migrating to the underside of the filter was counted by light microscopy 14 hours later (Figure 5B). Supernatants from T-bet^+^ memory B cells induced large numbers of BMDMs to cross transwell filters relative to all other conditions, where only the supernatants from classical memory B cells showed any capacity to induce migration above media baseline (Figure 5C).

**Figure 5.**
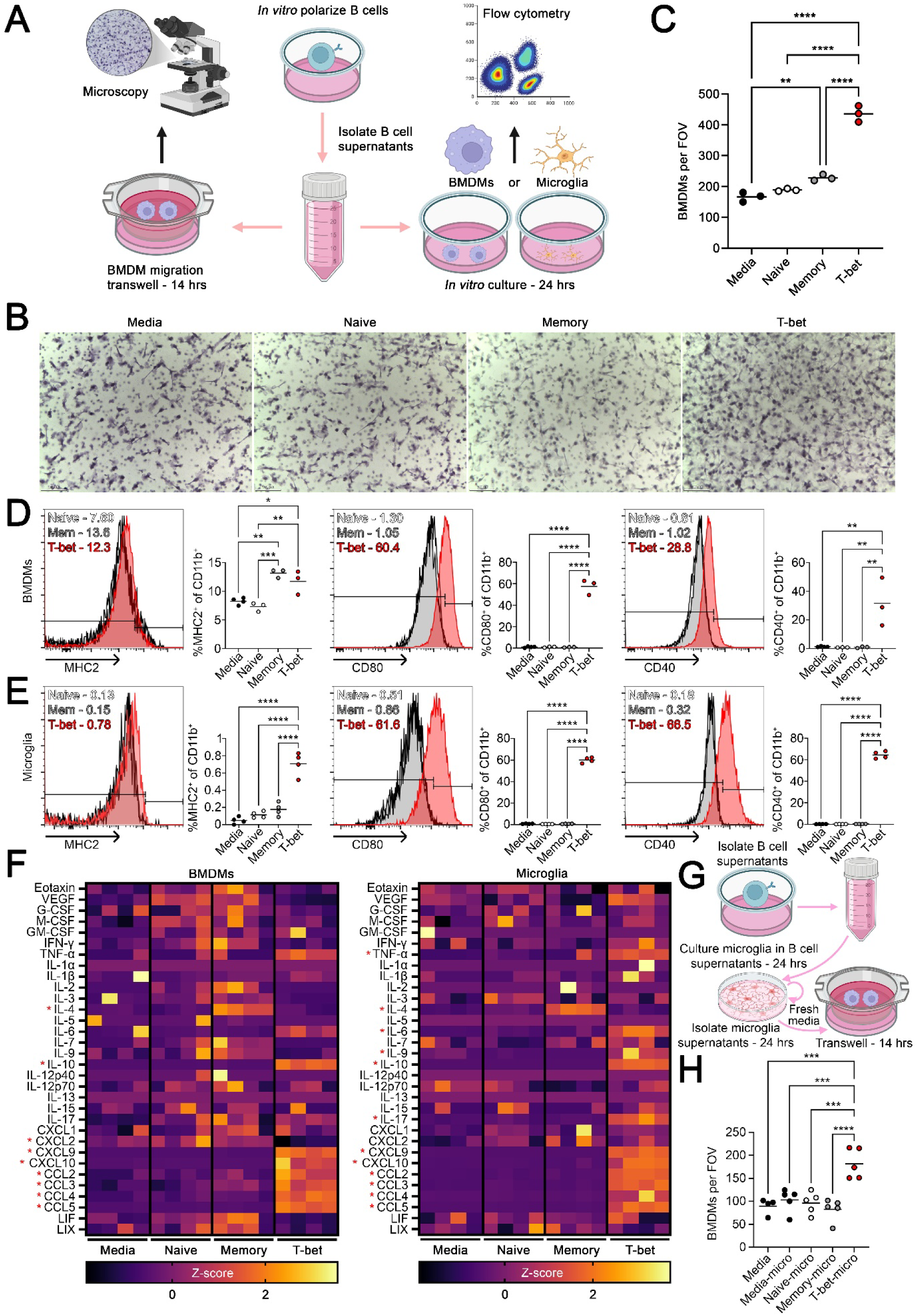
Secreted factors from T-bet^+^ memory B cells orchestrate microglia and macrophage activation and macrophage migration. (A) Schematic of *in vitro* experiments. Supernatants from *in vitro* polarized B cells are used in transwell assays (left) by placing supernatants below transwell filters and placing BMDMs on top. Fourteen hours later migration across the transwells is assessed by microscopy. Alternatively, supernatants are placed on BMDMs or microglia for 24 hours (right) then the expression of surface markers on BMDMs or microglia is assessed by flow cytometry. (B) Representative images of transwells quantified in C (n = 3 technical replicates). Scale bars are 100 μm. Representative 1 of 5 experiments. (D and E) Quantification and histograms showing MHC2, CD80, and CD40 expression on BMDMs (D) and microglia (E) as assessed by flow cytometry (n = 3-4 technical replicates). (D) Representative 1 of 4 experiments. (E) Representative 1 of 3 experiments. (F) BMDMs or microglia were incubated with the indicated B cell supernatants for 24 hours then incubated in blank media for 24 additional hours. These supernatants from BMDMs or microglia were then analyzed using a Luminex and summarized as Z-scored expression of each factor in a heatmap (n = 4). Representative 1 of 2 experiments. (G) Schematic demonstrating the usage of B cell stimulated microglial supernatants in transwell assays to assess their capacity to induce BMDM migration over 7 hours that is quantified in H (n = 5 technical replicates). Representative 1 of 3 experiments. Statistical significance was assessed using a parametric one-way ANOVA with a tukey’s post-hoc test (C, D, E, F, and H). Data are presented as mean (C, D, E, and H) or individual values in a heatmap (F) and p-values are summarized as *p<0.05, **p<0.01, ***p<0.001, and ****p<0.0001. See also Figure S5.

In parallel, we determined that supernatants from T-bet^+^ memory B cells, but not other B cells, potently induced CD80 and CD40 expression on both microglia and BMDMs after a 24 hour incubation (Figure 5D and 5E). Supernatants from T-bet^+^ memory B cells had a small effect on MHC2 expression by microglia and BMDMs, with classical memory B cell supernatants also having a similar outcome. Next, we switched the B cell supernatants that were on microglia or BMDMs after 24 hours of incubation with fresh culture media for 24 hours, then collected these supernatants for analysis using Luminex. Supernatants from T-bet^+^ memory B cells elicited IL-10, CXCL9, CXCL10, CCL2, CCL3, CCL4, and CCL5 production from both BMDMs and microglia suggesting they generally induced the expression of a range of chemokines (Figure 5F and Supplemental 5A). Microglia also produced more TNFα, IL-6, IL-9, and IL-17 in response to T-bet^+^ memory B cell supernatants. Analysis of the stimulated BMDMs and microglia using microscopy demonstrated that BMDMs and microglia switched from a spindle-like morphology to a flattened form (Supplemental figure 5B and 5C). Additionally, their numbers and expression level of Iba1 increased. Thus, T-bet^+^ memory B cell supernatants are potent activators of both macrophages and microglia.

An anomaly was the observation that the transfer of T-bet^+^ memory B cells promoted macrophages to enter the CNS parenchyma, yet, T-bet^+^ memory B cells were mostly confined within the meninges and perivascular cuffs. As T-bet^+^ memory B cells induced chemokine expression by microglia (Figure 5F and Supplemental 5A), we hypothesized that microglial chemokines in response to T-bet^+^ memory B cell exposure promote macrophage migration and accumulation within the CNS parenchyma. To test this, we stimulated microglia with supernatants from B cell cultures for 24 hours, then removed the media and incubated the microglia in fresh media for 24 more hours (Figure 5G). The latter supernatants were then tested. Only supernatants from microglia stimulated with T-bet^+^ memory B cell supernatants enhanced BMDM migration across transwell filters above a media-only control (Figure 5H). Thus, in addition to T-bet^+^ memory B cells inducing macrophage migration, microglia stimulated by T-bet^+^ memory B cells also facilitated macrophage migration.

### T-bet^+^ memory B cells have an immune profile characterized by inflammatory chemokine and cytokine expression

To determine mechanistically how T-bet^+^ memory B cells were stimulating microglia and macrophage, we polarized B cells *in vitro* for four days, then analyzed the B cells using bulk RNA sequencing. In a parallel experiment, B cells were polarized for three days then switched into culture media containing only BAFF protein and these supernatants were collected for Luminex analysis after 24 hours. By Luminex, T-bet^+^ memory B cells expressed large amounts of VEGF, G-CSF, GM-CSF, IFN-γ, TNF-α, IL-3, IL-6, IL-9, IL-10, IL-15, IL-17, CXCL1, CXCL9, CXCL10, CCL3, CCL4, CCL5, and LIF (Figure 6A and Supplemental figure 6A). Bulk RNA sequencing (Supplemental figure 6B and 6C and Supplemental table 2) showed elevation in transcripts of several of these molecules although many of these trended towards significance. Thus, T-bet^+^ memory B cells upregulate several inflammatory cytokines and chemokines that could explain their pathogenic potential.

**Figure 6.**
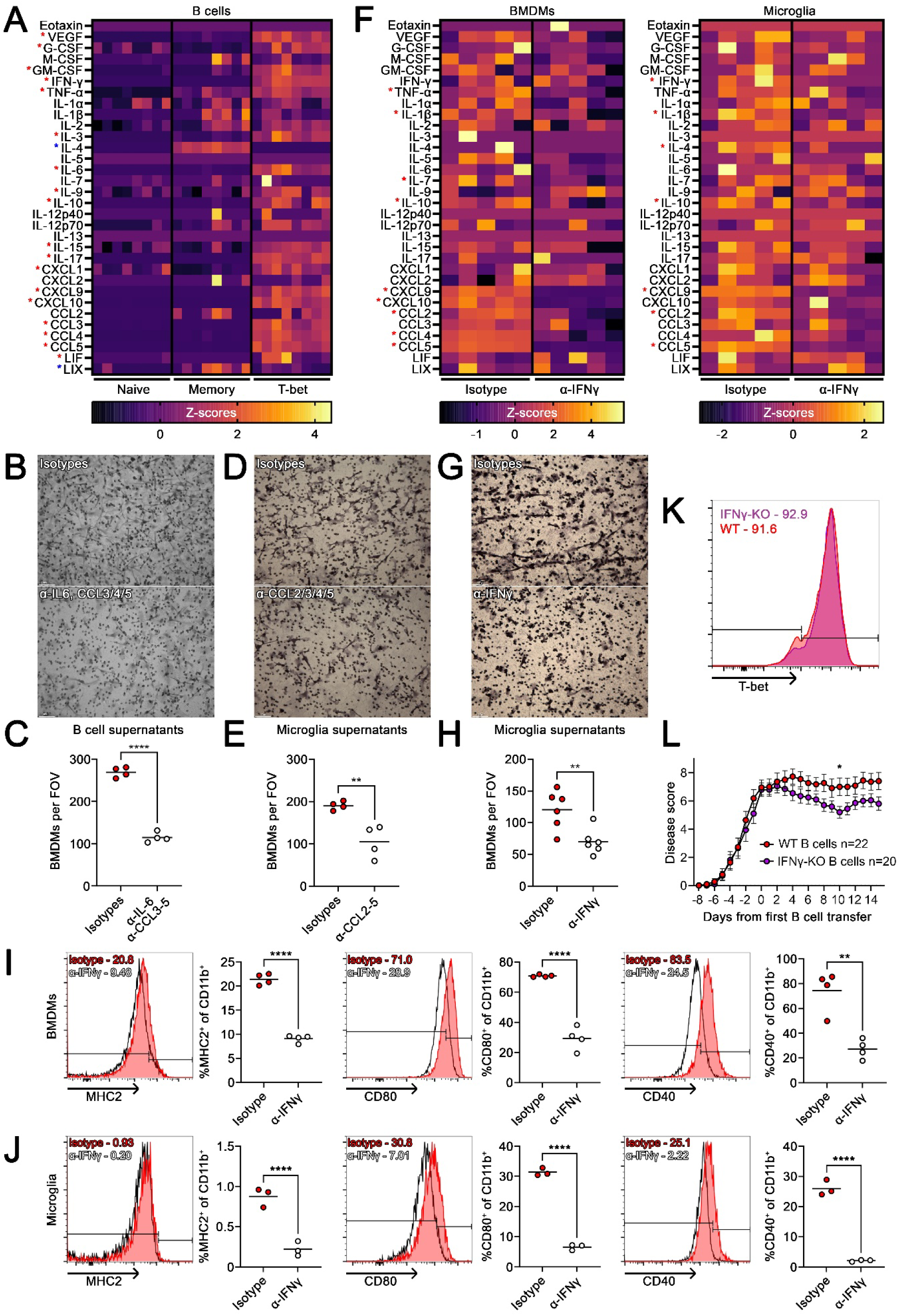
Chemokines, IL-6, and IFN-γ from T-bet^+^ memory B cells orchestrate myeloid cell activation and migration. (A) B cells were *in vitro* polarized into the indicated B cell types for 3 days then switched into blank media containing only BAFF survival factor for 24 hours. These supernatants were then analyzed using a Luminex and the relative expression of the indicated proteins is displayed using Z-scored values in a heatmap (n = 8). Red significance indicates a difference between the T-bet group and others and blue significance indicates a difference between memory B cells and others. Combination of two independent experiments. (B and C) Supernatants from T-bet^+^ memory B cells were incubated with isotype antibodies or antibodies against IL-6, CCL3, CCL4, and CCL5 for half an hour then tested for their capacity to induce BMDM migration over 7 hours (n = 4 technical replicates). Representative 1 of 4 independent experiments. Scale bars are 100 μm. (D and E) Supernatants from microglia stimulated with T-bet^+^ memory B cell supernatants were incubated with isotype antibodies or antibodies against CCL2, CCL3, CCL4, and CCL5 for half an hour then tested for their capacity to induce BMDM migration over 7 hours (n = 4 technical replicates). Representative 1 of 3 independent experiments. Scale bars are 100 μm. (F) Supernatants from T-bet^+^ memory B cells were incubated with an isotype antibody or anti-IFNγ for half an hour then used to stimulate BMDMs or microglia for 24 hours. BMDMs and microglia were then incubated in blank media for 24 hours and these supernatants were analyzed using a Luminex and presented as Z-scored heatmaps (n = 5). Representative 1 of 2 independent experiments. (G and H) The supernatants generated from microglia as described in F were tested for their capacity to induce BMDM migration using a transwell over 7 hours (n = 6 technical replicates). Representative 1 of 3 independent experiments. Scale bars are 100 μm. (I and J) Supernatants from T-bet^+^ memory B cells were incubated with an isotype antibody or anti-IFNγ for half an hour then used to stimulate BMDMs (top) or microglia (bottom) for 24 hours. Expression of MHC2, CD80, and CD40 were assessed using flow cytometry as shown though histograms and quantification (n = 3-4 technical replicates). Representative 1 of 3 independent experiments. (K) Histogram showing T-bet expression in B cells from wildtype or IFNγ-KO mice that were polarized *in vitro* into T-bet^+^ memory B cells. (L) EAE diseases scores showing MOG_35-55_ induced EAE in wildtype mice where *in vitro* polarized T-bet^+^ memory B cells from wildtype or IFNγ-KO mice were transferred at peak disease and 7 days later, then tracked for an additional 8 days after second transfer. Combination of 3 independent experiments (n = 20-22). Statistical significance was assessed using a parametric one-way ANOVA with a tukey’s post-hoc test (A), a student’s T-test (C, E, F, H, I, and J), or multiple Mann-Whitney U tests (L). Data are presented as mean (C, E, H, I, and J), mean ± SEM (L), or individual values (A and F) and p-values are summarized as *p<0.05, **p<0.01, and ****p<0.0001. See also Figure S6.

By incorporating neutralizing antibodies into the transwell assay, we determined that blocking IL-6, CCL3, CCL4 and CCL5 in T-bet^+^ memory B cell supernatants attenuated their capacity to induce BMDM migration (Figure 6B and 6C). Moreover, microglia exposed to T-bet^+^ memory B cell supernatants also induced BMDM migration through CCL2, CCL3, CCL4 and CCL5 (Figure 6D and 6E). Thus, T-bet^+^ memory B cells induce macrophage migration directly, and indirectly through microglia, using IL-6, CCL2, CCL3, CCL4, and CCL5.

### IFN-γ is critical for T-bet^+^ memory B cell pathogenicity

As large amounts of IFN-γ were detected in the supernatants isolated from T-bet^+^ memory B cells (Figure 6A and Supplemental figure 6C), we evaluated whether IFN-γ in T-bet^+^ memory B cell supernatants was responsible for inducing macrophage/microglia activation. Supernatants from T-bet^+^ memory B cells were incubated with either isotype or anti-IFN-γ antibody and then used to stimulate BMDMs or microglia. After 24 hours, these supernatants were removed and new media was added to the BMDMs or microglia and incubated for another day, then analyzed using a Luminex assay. Blocking IFN-γ reduced IL-1β, IL-10, CXCL9, CCL2, and CCL5 in both BMDMs and microglia (Figure 6F). Additional reductions of TNF-α, IL-7, CXCL10, and CCL4 were observed for BMDMs with similar trends observed for some of the chemokines in microglia. Microglia had additional reductions in IFN-γ and IL-4. Thus, a major T-bet^+^ memory B cell factor that induced cytokines/chemokines in microglia and BMDMs was IFN-γ.

Next, supernatants from microglia that were stimulated with T-bet^+^ memory B cell supernatants, with or without IFN-γ neutralization, were tested. We found that IFN-γ neutralization in T-bet^+^ memory B cells supernatants reduced the capacity of microglia to induce BMDM migration across transwells (Figure 6G and 6H).

We compared the capacity of T-bet^+^ memory B cell supernatants, with isotype or IFN-γ neutralizing antibody, to promote MHC2, CD80, and CD40 expression on BMDMs and microglia. For both myeloid cell types, neutralization of IFN-γ reduced the expression of each of these markers (Figure 6I and 6J). Thus, IFN-γ from T-bet^+^ memory B cells was critical for the induction of inflammatory polarization of macrophages and microglia.

Finally, we addressed whether disrupting IFN-γ in T-bet^+^ memory B cells would affect their capacity to impact EAE. We *in vitro* polarized B cells from wild type or IFN-γ-KO^61^ mice into T-bet^+^ memory B cells, finding no difference in their capacity to form T-bet^+^ memory B cells (Figure 6K). Cells were transferred into MOG_35-55_ EAE mice at peak disease and 7 days later, then evaluated for their EAE disability and immune cell infiltration into the spinal cord by flow cytometry. While the EAE scores started identically, mice receiving IFN-γ-KO T-bet^+^ memory B cells began to recover compared to those receiving wildtype T-bet^+^ memory B cells, resulting in a significant difference 10 days after the first cell transfer (Figure 6L). After this initial recovery, however, the IFN-γ-KO T-bet^+^ memory B cell transferred group had a slight worsening thereby diminishing the statistical difference between groups. Analysis of the spinal cords at this later period revealed no difference in CD4^+^ T cell, B cell, neutrophil, microglia or macrophage numbers (Supplemental figure 6D). Unsupervised clustering of microglia revealed no changes in microglia states and similar analysis on macrophages revealed only a small change by proportion in macrophage states (Supplemental figure 6E-6J), a slight increase in Ly-6C^hi^ CD115^hi^ monocytes in mice receiving wildtype T-bet^+^ memory B cells, but no change by absolute numbers. Thus, by the end of the experiment both groups had nearly identical immune cell infiltrates.

### T-bet expression in B cells affects MOG_35-55_ induced EAE severity in an age-dependent manner

The proportion of B cells expressing T-bet increases with age^34^, which we confirmed (Supplemental figure 7A and 7B). Thus, the accumulation of these cells with age could promote EAE severity. To test this, we crossed Mb1-cre with Tbx21^flox/flox^ mice to generate mice with wild type T-bet expression (Mb1-cre^-^Tbx21^flox/flox^, WT) or with a B cell specific knock-out of T-bet (Mb1-cre^+^ Tbx21^flox/flox^, KO). These mice were then aged to 80-92 weeks of age (old) or 8-16 weeks of age (young). Deletion of T-bet in aged B cells, but not in CD4^+^ T cells, was confirmed in unimmunized animals (Figure 7A) and was corroborated by polarizing B cells from T-bet knock-out mice into T-bet^+^ memory cells (Supplemental figure 7C). EAE was then induced in young and aged mice using MOG_35-55_. In young animals, there was no difference in EAE severity between genotypes consistent with the low numbers of T-bet^+^ memory B cells found in wild type young mice, and with the knowledge that MOG_35-55_ EAE is B cell-independent^45–47^. Conversely, there was a large difference in disease severity in aged animals where removing T-bet in B cells greatly reduced EAE severity (Figure 7B). Thus, with aging, the severity of MOG_35-55_ EAE becomes highly dependent on T-bet^+^ memory cells.

**Figure 7.**
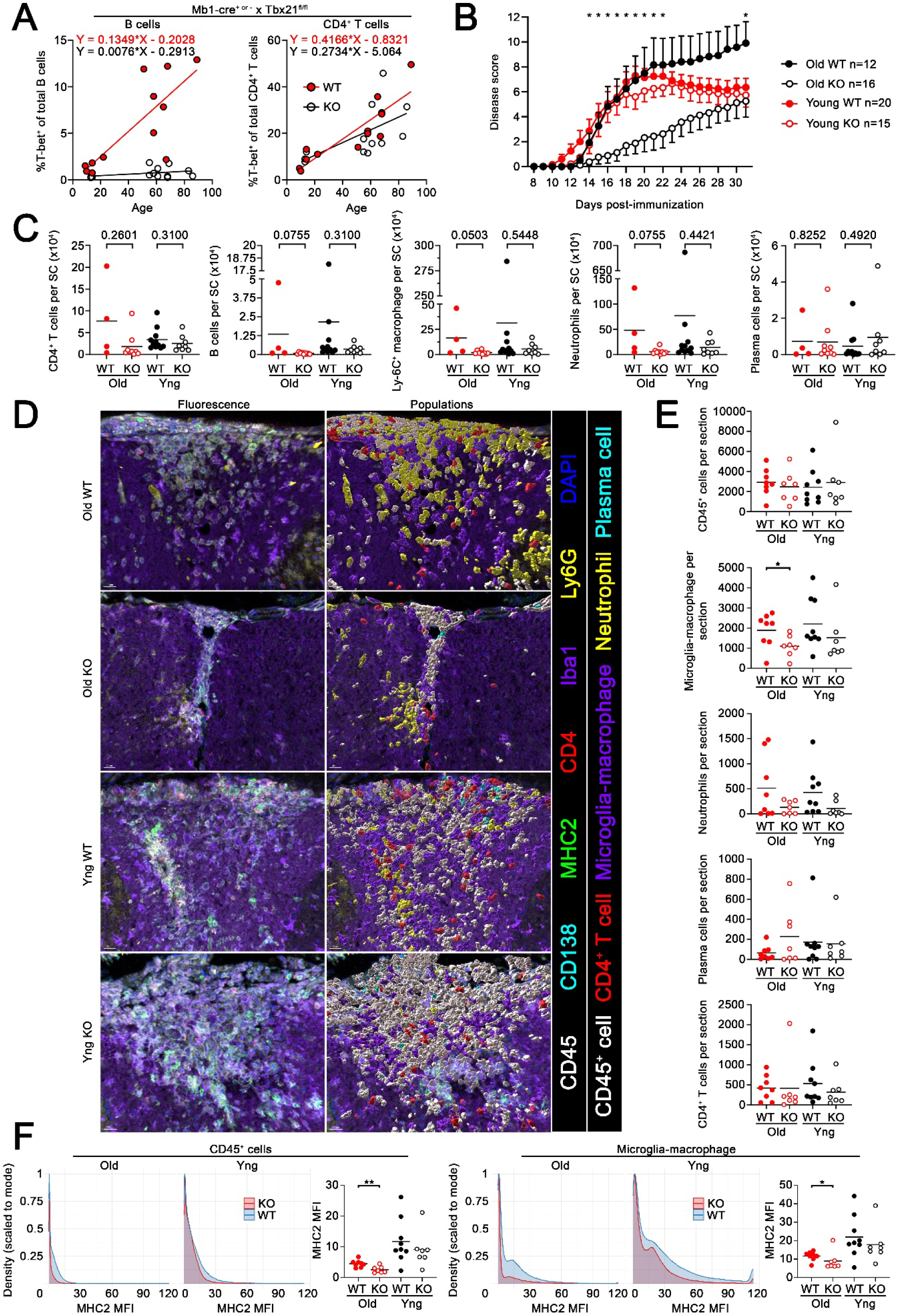
T-bet expression in B cells affects MOG_35-55_ induced EAE severity in an age-dependent manner. (A) T-bet expression in CD19^+^ B220^+^ B cells (left) or CD4^+^ T cells (right) in the spleens of unimmunized Mb1-cre^+/-or-/-^ Tbx21^fl/fl^ mice at various ages (weeks). Linear regressions are shown in red for Mb1-cre^-/-^ mice and black from Mb1-cre^+/-^ mice. (B-F) Mb1-cre^+/-^ Tbx21^fl/fl^ mice (KO) or Mb1-cre^-/-^ Tbx21^fl/fl^ mice (WT) that were young (yng, 7-16 weeks of age) or old (81 to 92 weeks of age) were immunized with MOG_35-55_ to induce EAE (n = 12-20). (B) EAE disease curve depicting disease severity over time. Represents the combination of 13 experiments. (C) Quantification of immune cell infiltrates into spinal cords of mice at termination of experiment (B) using flow cytometry (n = 4-11). (D) Representative images of CD45, CD138, MHC2, CD4, Iba1, Ly6G, and DAPI staining in the spinal cords of mice (left) and the populations identified through histoflow cytometry (right); mice were from the termination of experiment of panel B. Scale bars are 20 μm. (E) Quantification of the absolute number of each indicated population per spinal cord section (n = 7-9). (F) Histograms show the mean fluorescence intensity of MHC2 on CD45^+^ cells (left) and macrophage-microglia (right) and quantification. Representative of one experiment. Statistical significance was assessed using single or multiple Mann-Whitney U tests comparing wildtype to knockout within each age bracket (B, C, E, and F). Data are presented as mean + or - SEM (B) or mean (C, E, and F) and p-values are summarized as *p<0.05 and **p<0.01, or explicitly stated. See also Figure S7.

Analysis by flow cytometry (Supplemental figure 7D) detected trends towards reduced numbers of CD4^+^ T cells, B cells, Ly-6C^+^ macrophages, and neutrophils in the CNS of aged B cell specific T-bet knock-out mice relative to aged wildtype mice (Figure 7C). In contrast, in young mice, no differences in immune cell infiltrates into the CNS were found between wild type or knock-out animals. Plasma cell numbers were not different for either age group. Histoflow cytometry (Supplemental figure 7E) using CD138, B220, MHC2, Ly6G, CD45, CD4, Iba1, and T-bet markers yielded 5 major populations: CD45^+^ cells, macrophage-microglia, neutrophils, plasma cells, and CD4^+^ T cells (Figure 7D and Supplemental figure 7F). B cells did not adequately resolve into a cluster sufficient for analysis, but they were observed in the CNS of each group where T-bet expression was only found in the CNS of wild type animals. T-bet expression in B220^+^ B cells was extremely rare in young mice but could be found in aged animals (Supplemental figure 7G) although most B cells were T-bet^-^. There were no changes in the numbers of immune cells in the spinal cords of young mice or in the expression of MHC2 on these cells suggesting immune cell infiltrates were identical between the groups (Figure 7E and 7F). Conversely, in the aged CNS there was an increase in the number of microglia/macrophages in wild type animals over knock-out aged mice but no statistically significant changes in other populations (Figure 7E). Additionally, there was increased expression of MHC2 on CD45^+^ cells and microglia/macrophages in old wild type mice suggesting that the activation profile of these cells decreased when T-bet expression was removed from B cells (Figure 7F). Overall, these data demonstrate that deletion of T-bet in B cells reduces EAE severity in an age dependent manner, and most potently by modifying microglia and macrophage numbers and activation.

## Discussion

Here we demonstrate that T-bet^+^ memory B cells found in MS lesions are pathogenic in EAE. Critically, we find that T-bet^+^ memory B cells coordinate the recruitment of monocytes/macrophages into the inflamed CNS and activate microglia and monocytes/macrophages to worsen EAE disability and demyelination. This appears to be achieved through IL-6, CCL3, CCL4, and CCL5 from T-bet^+^ memory B cells that activate and recruit macrophages; and by IFN-γ from T-bet^+^ memory B cells that further induce migration and activation of macrophages. Further pathogenicity of T-bet^+^ memory B cells is found in aging, a factor that elevates T-bet expressing B cells^34,62^, where aging mice with T-bet deleted in B cells had attenuated EAE severity and myeloid accumulation compared to their wildtype controls.

A similar population of B cells has also been observed in MS that is characterized by high expression of T-bet, IFN-γ, IL-6, CCL3, CCL4, CCL5, and many of the other inflammatory proteins that we observed to be associated with T-bet^+^ memory B cells^63^. While pathogenicity was not previously demonstrated, this population was associated with the conversion of clinically isolated syndrome to MS, and to be closely associated with Epstein-Barr Virus (EBV) infection that is linked to the development of MS^64^. Indeed, EBV infection is known to induce T-bet expression in B cells^65^. Experiments using gamma-herpes virus infection of EAE mice, modelling EBV infection, have found that deletion of T-bet in B cells attenuates the capacity of gamma-herpes viruses to worsen EAE severity^66^. These results mirror what we found in aging mice where T-bet deletion attenuated MOG_35-55_ induced EAE. Currently it is unclear whether aging and EBV-infection would promote T-bet^+^ memory B cell activity through similar or different mechanisms.

One possibility is that T-bet^+^ memory B cells can be pathogenic regardless of how they were generated. We found that T-bet^+^ memory B cells can be pathogenic independent of autoreactivity, as the transfer of wildtype and MD4 T-bet^+^ memory B cells did not differ in their capacity to worsen EAE, making such a scenario possible. However, we did not test whether autoreactivity could amplify the pathogenicity of T-bet^+^ memory B cells which is a distinct possibility given that others have demonstrated that these cells are likely to be autoreactive in MS^67^, can participate in cognate interactions with T cells^67,68^, and can be induced to produce autoreactive antibodies^67,69^.

Others have described interactions between B cells and macrophage/microglia in the context of MS. This literature includes B cells in MS promoting inflammatory polarization of macrophages through GM-CSF production^15^, and soluble factors from MS B cells inducing the expression of CD80 on microglia and macrophages^70^. Conversely, others have shown that IL-10 production by MS B cells can suppress neuroinflammation^71^. Adding to this, we found that T-bet^+^ memory B cells promoted an overall inflammatory polarization in macrophage and microglia primarily through IFN-γ; however, we observed that they also produce IL-10. These two cytokines are known to polarize macrophage in different ways, IFN-γ promoting an inflammatory polarization while IL-10 facilitates a regulatory state^72^; moreover, they are known to act antagonistic to one another^73–76^. Our *in vitro* experiments in combination with our *in vivo* data suggest that the effects of IFN-γ overtake those of IL-10 in wild type T-bet^+^ memory B cells, as we observed increases in CD40 and CD80 expression, known to be affected by IFN-γ^77^, suggesting potent IFN-γ stimulation. In our experiments using IFN-γ KO T-bet^+^ memory B cells, we observed reduced severity of EAE during the recovery phase of EAE but no effect on whether secondary worsening occurred. These results suggest that the role of T-bet^+^ memory B cell derived IFN-γ *in vivo* is not to promote secondary waves of inflammation, but rather that it antagonizes the activity of IL-10 to maintain inflammation and delay recovery. The lack of an effect outside of the recovery phase could potentially be related to T-cells also producing IFN-γ^78^ during EAE where T-bet^+^ memory B cells may be redundant during some stages of inflammation.

Our results have demonstrated that T-bet^+^ memory B cells delivered into the blood will enter the CNS and contribute to CNS pathology if neuroinflammation is already present. This would suggest that anti-CD20 monoclonal antibodies that robustly deplete peripheral B cells^79^ would work in part through the depletion of these cells peripherally. We have demonstrated that T-bet^+^ memory B cells are found in MS lesions consistent with the findings of others that atypical memory B cells concentrate in CSF^42^, CXCR3^+^ B cells are concentrated in the CNS^43,63^, and are in the meninges of MS patients^67^. Currently it is not clear whether anti-CD20 monoclonal antibodies effectively clear B cells that are in CNS barriers^80,81^. Indeed, once compartmentalized inflammation is initiated it is possible that B cells could self-renew^82^ maintaining a T-bet^+^ memory B cell population. In this case, CNS-penetrant drugs such as Bruton’s tyrosine kinase inhibitors^83^ may be useful to suppress their activity as BTK signaling is required for T-bet^+^ memory B cells to maintain their phenotype^84,85^.

Overall, we provide new insights that T-bet^+^ memory B cells are pathogenic in EAE, and by inference, MS, and that their capacity to modify the macrophage and microglial immune response is critical to their pathogenicity.

### Limitations of the study

The experiments described in this work all use female animals and thus, future work will need to determine whether sex differences impact the pathogenicity of T-bet^+^ memory B cells.

Additionally, the experimental work was completed in mice. Human T-bet^+^ memory B cells have been described to make the same implicated proteins described here^63^ but future studies will need to confirm that human T-bet^+^ memory B cells similarly affect human microglia and macrophage.

Due to the limited number of MS lesions analyzed, we could not determine whether there were changes in the proportions of these cells in each lesion type, between sexes, or whether they differ with age. Larger datasets will be needed to determine if these factors affect T-bet^+^ memory B cell accumulation.

## Resource availability

### Lead contact

Any requests for resources should be directed to the lead contact, V. Wee Yong, who will fulfill all requests.

## Materials availability

No new reagents were produced in this study. All the reagents used in this study are as described in the key resources table.

## Data availability

Bulk RNA sequencing data generated in this study will be made publicly available as of the date of publication on the NCBI Sequence Read Archive (PRJNA1480860). Code is available upon request. Any additional information required to reanalyze the data reported in this paper is available from the lead contact upon request.

## Acknowledgements

This study was supported by a Foundation grant to V.W.Y. from the Canadian Institutes of Health Research. R.W.J acknowledges postdoctoral fellowship funding from the University of Calgary Eyes High program, Multiple Sclerosis Canada, and a Roche unrestricted educational fellowship. We thank the Hotchkiss Brain Institute Advanced Microscopy Platform for imaging infrastructure and expertise. We thank Dr. Alexandre Rudensky for providing us with the Tbx21^tdTomato-T2A-creERT2^ mouse strain. We thank the UK Multiple Sclerosis and Parkinson’s Tissue Bank at Imperial College, London, and Dr. Djordje Gveric for providing non-neurological control and MS brain tissues.

## Author contributions

R.W.J. and V.W.Y. conceived and designed the study. R.W.J. conducted the experiments and supervised the analysis of the experiments. M.G. conducted bioinformatic analysis on bulk RNA sequencing data. M.G., A.M., S.X., and S.Z. contributed to the cell counting of the transwell assays. S.X. contributed to the demyelination image analysis. A.M. contributed to tissue staining and imaging of tissues. R.G. characterized the MS lesions and cut tissue sections. M.T.M. contributed to the Luminex assays. A.P. contributed some of the MS tissues. R.L. provided helpful discussions and B cell expertise. R.W.J. and V.W.Y. wrote the manuscript with contributions from the other authors.

## Declaration of interests

The authors declare no competing interests.

## Declaration of generative AI and AI-assisted technologies in the writing process

The authors did not make use of AI during the writing process.

## Methods

### Animals and EAE induction

Female C57BL/6 mice were purchased from Jackson Laboratories and used in experiments within 7-16 weeks of age. Mice were immunized with MOG35-55 peptide (50 μg/100 μL) emulsified in CFA supplemented with 5 mg/mL heat-inactivated *Mycobacterium tuberculosis* H37Ra; 50 μL emulsion was deposited on either side of the tail base. Pertussis toxin (PTX) (300 ng/200 μL) was injected i.p. on days 0 and 2 after MOG immunization. Mice were scored daily using a 15-point scale described previously^86^. Mice were housed in a specific pathogen-free facility. All animals were handled in accordance with guidelines of the Canadian Council for Animal Care and their use was approved by the animal ethics committee of the University of Calgary.

### B cell isolation and polarization

The spleen and inguinal, axial, and brachial lymph nodes were isolated from naïve mice and dissociated using frosted glass slides in ice cold easysep buffer (1mM EDTA, 2% FBS in PBS). B cells were isolated from the dissociated cells using an EasySep Negative selection Mouse B cell Enrichment Kit according to the manufacturers protocol.

Isolated B cells were then cultured in B cell media (RPMI with L-glutamine) supplemented with 10% FBS, 100 U/mL penicillin-streptomycin, 1x GlutaMAX, 1x MEM non-essential amino acids solution, and 50 µM β-Mercaptoethanol at a concentration of 3 million B cells per mL at 37°C 5% CO_2_. B cells were cultured in this media for 4 days, with media changes on days 2 and 3, with the following supplements to polarize B cells towards distinct cell types: BAFF protein (50 ng/ml), ODN1828 CpG nucleotides (1.2 µg/mL), murine IFN-γ protein (50 ng/ml), Goat anti-mouse IgM/IgG (2.4 µg/mL), murine IL-4 protein (100 ng/ml), and rat anti-mouse CD40 (5 µg/mL). Naïve B cells were generated by culturing isolated B cells with BAFF. Memory B cells were generated by culturing isolated B cells with BAFF, murine IL-4 protein, Goat anti-mouse IgM/IgG, and rat anti-mouse CD40. T-bet^+^ memory B cells were generated by culturing isolated B cells with BAFF, murine IFN-γ protein, ODN1828 CpG nucleotides, Goat anti-mouse IgM/IgG, and rat anti-mouse CD40.

For *in vivo* experiments using *in vitro* polarized B cells, the B cells were isolated by centrifuging them at 500 x g for 10 minutes at 4°C after a total of 4 days in culture. Cells were then washed with 40 mL of ice cold HBSS (no calcium, magnesium, or phenol red), then centrifuged again. B cells were then suspended in ice cold HBSS again at a concentration of 40-50 million B cells per mL and kept on ice until 200 μL were injected intravenously into mice. In all experiments, groups were injected with equal numbers of live B cells as determined by trypan blue staining and counting using a hemocytometer. The viability and numbers of B cells were determined using trypan blue staining and a hemocytometer. Cells were injected into EAE mice at peak disease and again 7 days later.

For *in vitro* experiments, B cells were either polarized as written above or were polarized for three days then on the third day the cells were centrifuged and suspended in 40 mL of ice cold HBSS. B cells were then centrifuged and washed again in ice cold HBSS before again being centrifuged. B cells were then suspended in complete RPMI with BAFF protein and cultured for 24 hours at 37°C 5% CO_2_.

### Multiple sclerosis samples

Postmortem frozen brain tissues from people with MS and healthy control brain tissue were obtained from The Multiple Sclerosis and Parkinson’s Tissue Bank situated at Imperial College, London. This bank has been approved as a Research Tissue Bank by the Wales Research Ethics Committee (Ref. No. 18/WA/0238). Additional MS tissues were obtained from Dr. Alex Prat, University of Montreal with full ethical approval (BH07.001, Nagano 20.332–YP) and informed consent as approved by the CRCHUM and University of Montreal research ethics committee. The use of these human tissues in Calgary for research was approved by the Conjoint Health Research Ethics Board at the University of Calgary (Ethics ID REB15-0444). Patient characteristics are summarized in Supplemental table 1.

MS sections were classified based on PLP and NeuN staining to define white and grey matter and HLA-DR to identify lesions with immune activity. White matter areas with patches of PLP loss were defined as active lesions if there were confluent HLA-DR^+^ cells throughout the demyelinated area or as mixed active/inactive lesions if HLA-DR^+^ cells were present only at the edge of the demyelinated area. Meningeal inflammation was defined as the accumulation of HLA-DR^+^ cells in the meninges.

### Analysis of MS samples

Low magnification (10x) images of the DAPI channel were taken to determine the tissue structure using a Leica stellaris 5. The structure was cross-referenced with the characterized lesion structures determined using HLA-DR, PLP1, Neun, and DAPI staining. Using a 20x magnification lens, all CD20^+^ cells in each anatomical region were imaged in the DAPI, CD20, and T-bet channels. Images in Lif files were converted to tif files and then ims files for analysis in Imaris. The number of CD20^+^ cells and the number of T-bet^+^ CD20^+^ cells were manually counted in each image. To calculate the percentage of B cells expressing T-bet in each anatomical region, the number of T-bet^+^ CD20^+^ cells in that area across all images were divided by the number of CD20^+^ cells in the same area across all images.

### Tissue preparation for histology

On the day of tissue collection, mice were given a sublethal dose of ketamine/xylazine and perfused with 10 mL of ice-cold phosphate-buffered saline (PBS) followed with 5 mL of ice-cold 4% paraformaldehyde (PFA) via injection into the left ventricle. Inguinal lymph nodes draining the site of immunization or spleens were then fixed overnight in 2% PFA at 4°C and protected from light, then transferred into 30% sucrose for 24 hours. Whole spinal columns were taken from mice and fixed overnight in 2% PFA then washed once in PBS. To decalcify the spinal columns, they were then transferred into 0.1M EDTA PBS, which was refreshed daily, for 4 days, during which the columns were kept at room temperature and protected from light with gentle rocking. Columns were then transferred into 30% sucrose for 2-7 days at 4°C. Tissues were then mounted in blocks of OCT compound and stored at −80°C until use. Tissues were either mounted longitudinally or coronally with the cervical to lumbar sections being used for histological analysis. Tissue blocks were cut into 14 μm thick sections and stored at −20°C until use. Fresh frozen human tissue sections were cut into 10 μm thick sections then frozen at −80°C until use.

### Immunofluorescence histology

Human and mouse tissues were briefly rehydrated in ice-cold PBS. Human sections were fixed in 2% PFA for 20 mins at room temperature and washed three times in PBS. The human tissues were then permeabilized in 0.2% Triton X-100 and 0.05% Tween-20 for 20 mins at room temperature. The slides were then washed 3 times before proceeding. For demyelination analysis, sections were delipidated by sequential wash with 50%, 70%, 90%, 95%, 100%, 95%, 90%, 70%, and 50% ethanol then washed in PBS three times before proceeding. Tissues were then blocked for 2 hours with 10% Horse serum, 1% BSA, 0.1% cold fish gelatin, 0.1% Triton X-100, and 0.05% Tween-20 in PBS. Tissues were then stained overnight at 4°C with nuclear yellow (1:3000-1:10000, 2 mg/mL) in staining buffer (1% BSA, 0.1% cold fish gelatin, and 0.5% Triton X-100 in PBS) with primary antibodies as listed in the key resources table. After overnight staining, slides were washed three times in PBS then stained with TrueBlack Lipofuscin Autofluorescence Quencher according to the manufacturer’s instructions for 2 min at room temperature. Tissues were washed in PBS four times then stained at 4°C for 2-6 hours in staining buffer (without Triton X-100 and Tween-20) with secondary antibodies. After this step, samples were washed in PBS four times and blocked using the above blocking buffer (without Triton X-100 and Tween-20) plus 5% rat serum for 1 hour at 4°C. Tissues were then stained with fluorophore conjugated primary antibodies and DAPI (if used, 1:1000) in staining buffer (without Triton X-100 and Tween-20) at 4°C overnight. Tissues were then washed four times in PBS then mounted using Fluoromount-G.

Images were taken using a Leica TCS SP8 or stellaris 5 laser confocal microscope with a 10x air (0.40 numerical aperture (NA)), 20x air (0.75 NA), or a 25x water (0.95 NA) objective. For SP8 imaging, fluorophores were stimulated sequentially with 405, 488, 552, and 640 nm lasers and two fluorescent parameters were captured with each laser through tunable filters and imaged using high sensitivity hybrid detectors. For the stellaris 5, fluorophores were stimulated with a 405 nm laser and a tunable white light laser and 2-3 fluorescent parameters were captured through tunable filters and imaged using high sensitivity hybrid detectors. The same laser power, gain, filter value settings were used between samples. Images were captured using bidirectional scanning, 1.5 µm step z-stacks, 580 nm pinhole, 1-5 line average or accumulation, and a resolution of 2048×2048 or 1028×1028 x, y pixels as 8, 12, or 16-bit data.

For demyelination analysis, 10x magnification was used to image longitudinal spinal cord sections of the cervical and thoracic regions that were used for analysis. For the histoflow cytometry analyses in Figures 2 and 5 the three largest spinal cord lesions (in longitudinal sections of the cervical and thoracic regions) based on CD68 staining were identified in low resolution (512×512, 10x) images and then the three lesions were imaged at a higher resolution (1028×1028, 25x) and these were used for analysis. The analyses of Figure 3 were similar to the above description except the 3 largest collections of B220 B cells were imaged. For the lymph node and spleen analyses, representative images of B cell follicles were captured. For Figure 4, coronal images were collected from 2-4 cervical and 2-4 lumbar sections (2048×2048 20x).

### Imaging of 96 well plates

After the supernatants were removed for Luminex analyses, 100 µL of 4% PFA was added to each well of BMDMs/microglia and the cells were fixed for 20 mins at room temperature. The PFA was then removed and 100 µL of PBS was added to each well and the plates were stored at 4°C in tin foil until usage. Cells were then washed again with PBS then 150 µL of 0.1% Triton X-100 in PBS was added to the cells at room temperature for 5 mins. Cells were then washed once in PBS and 150 µL of Intercept blocking buffer was added to the wells and blocked for 1 hour at room temperature. Blocking buffer was removed and 100 µL of intercept antibody diluent with (chicken anti-Iba1 (1:500) and DAPI (1:1000)) was added to the cells and incubated at 4°C overnight. The next day the antibody diluent was removed, and the wells were washed twice with 150 µL of PBS. Then 150 µL of intercept antibody diluent with donkey anti-chicken A488 (1:300) was added to each well and incubated at room temperature for 2 hrs. The antibody diluent was then removed, and the wells were washed and imaged on an imageXpress Micro XLS High-Content Analysis System using a 20x (0.45 NA) lens. For each well, 9 separate fields of view were collected.

Data was analyzed using Olympus Cellsens Dimension using the same thresholding across samples to define cells based on DAPI and Iba1 staining that was used for MFI measurement, cell counts, and elliptical form factor analysis. For all measurements, values across all 9 fields of view were combined and averaged.

### Histoflow cytometry

Histoflow cytometry analysis was conducted as previously described^48,49^ with additional modifications. Briefly, spectral compensation was applied to images using Leica Application Suite X software based on a compensation matrix manually generated using single-colour stained images. Compensated images were then imported into Ilastik^87^ and pixel classifiers were trained on 4-5 images to recognize nuclei and cell bodies. Probability maps classifying the probability of pixels being cells/nuclei were then exported as HDF5 files and converted into tiff files. These tiff files were then converted into Imaris files along with the original images using ImarisFileConverter. These images were then combined into one Imaris file and the cell module was used to identify individual cells in the image by using the cell probability map to identify all the voxels containing cells and then splitting them such that each cell only had one nucleus as determined by the nuclei probability map. A standard threshold was used to identify nuclei and cells. In some cases, areas in the image would be identified by manually tracing the area in Imaris, sometimes aided by thresholding a channel such as laminin staining. Then a new channel would be created and the fluorescence values within the traced spaces would be set to the max value, and spaces outside would be set to 0.

Next, the mean fluorescence intensity of all channels in the cell or just within the nucleus was exported from Imaris in csv files along with unique cell identifiers. A python script was then used to collate all the fluorescence values for all the cells into a single csv file. These csv files were then converted into fcs files using FlowJo for analysis. Gating strategies were used to define populations as depicted throughout the manuscript. Values from the gated populations were then exported from FlowJo into csv files containing all the fluorescent values and the cell ID of all cells in that gate for one sample. A python script was then used to extract all the cell IDs that were inputted back into Imaris to identify where the cells in the gate appeared in the image. Gates were adjusted until the mapping was accurate across multiple samples. Once all gates were validated, the absolute cell numbers were exported from FlowJo and analyzed in excel.

Cells from 2-3 lesions were added together and averaged for the analyses in Figures 2 and 3. In Figure 4 cells from 3 lesions (with 2 replicates per lesion) were added together and averaged. For Figure 5, the cells from 2-4 coronal sections were added together and averaged for the cervical and thoracic portions of the spinal cord separately, then the values of the cervical and thoracic portions of the spinal cord were averaged together.

For Figures 4 and 5 cells exported from FlowJo were additionally analyzed in R prior to calculating cell abundances. For Figure 4, inter-sample variability was normalized based on the MFI DAPI signal such that the mean DAPI signal for each sample was calculated, then a scaling factor relative to the median DAPI MFI was determined. The MFI of all other markers corresponding to stained proteins and DAPI was then multiplied by the scaling factor to standardize the signals (all average DAPI MFIs being identical after scaling). Principal component analysis was then conducted on the scaled data and Harmony^88^ was used to correct for batch effects. The data was then clustered using Louvain clustering using the MFI of CD138, B220, MHC2, Ly-6G, CD45, CD4, Iba1, nuclear T-bet, Cell score (based on the Ilastik output). Clusters were then mapped back to the tissue and all clusters that mapped to cells in the tissue were reclustered as above but including the DAPI MFI, the cell volume, and two additional measures: the first was generated in Imaris where all voxels with a B220 MFI above 6 were set to a 65535 in a new channel, and another new channel that set all voxels with a B220 value of 6 and a DAPI signal of 400 as 65535. Cells from this reclustering were again mapped to tissue to identify clusters mapping to cells and were carried to another round of reclustering. These cells were again mapped to the tissues and clusters contaminated with debris were subclustered and mapped to the tissue to further remove debris. Uncontaminated clusters were then combined and reclustered to generate the final UMAP. Clusters were mapped back to the tissue and were annotated as microglia/macrophage, CD4^+^ T cells, neutrophils, plasma cells, or CD45^+^ cells. Cell abundances in clusters and mean fluorescence intensity of the markers per cluster were then exported for analysis in excel.

For Figure 5, fluorescence values were transformed using an inverse hyperbolic sine (arcsinh) function, then unsupervised clustering was performed on the transformed data using FlowSOM^89^ using six markers (CD68, MHC2, Dectin-1, Arg1, Iba1, and Nuclear Ki67) to generate 20 unique clusters. FlowSOM clusters were then combined based on hierarchical agglomerative clustering (Ward’s D2 linkage) on cluster-median marker expression profiles to combine clusters with similar profiles. Combined clusters were then mapped back to the tissue to ensure their properties matched what was found in the tissues. A UMAP was generated based on the FlowSOM clustering and then cells within lesions were separated from non-lesion cells using a 50000 threshold based on the tracing of the lesions done in Imaris (where a value of 65535 was set for all voxels in the lesion). Cell abundances in clusters and median fluorescence intensity of the markers per cluster were then exported for analysis in excel.

### Demyelination analysis

Images were analyzed using FIJI^90^. Z-stacks were flattened using a Z-projection of the max intensity. The ROI manager tool was used to create an outline of the parenchyma of the spinal cord sections based on GFAP staining then crop the image to only contain the CNS parenchyma. Separate outlines were used to identify grey matter in the images (non-lesion MBP low areas) and these areas were cropped out of the image, leaving only white matter. The areas for each of these areas were collected along with the distance per pixel. Images containing only white matter were then imported into Ilastik. A pixel classifier was trained on 5 images from each analyzed experiment separately to identify myelinated and demyelinated areas based on MBP staining. Once trained, the classifier was run on all images in the same experiment and the mask containing the pixels predicted to be myelinated were exported as HDF5 files and imported back into FIJI. The distance per pixel was adjusted back to the original value for the sample, then the area covered by myelinated pixels was determined using thresholding where the same value was used across all images in the same experiment. Finally, the percentage demyelination was calculated by dividing the total myelinated area by the total parenchyma area minus the grey matter area. Two to three spinal cord sections (cervical to thoracic) were imaged per mouse and the areas for these cords were added together for the calculation.

### Flow cytometry

On the day of tissue collection, mice were given a sublethal dose of ketamine/xylazine and perfused with 10 mL of ice-cold PBS via injection into the left ventricle. Blood was extracted from the right ventricle using a 22-gauge needle preloaded with 0.5M EDTA prior to perfusion. Blood was then centrifuged at 1200 rpm for 8 minutes and the plasma was collected, and the pellet was incubated with 1ml ACK lysis buffer for 5 minutes. FACS buffer (2% FBS-PBS) was then added to the samples and centrifuged again. Blood was then incubated with ACK lysis buffer once more and centrifuged.

Lymph nodes and spleens were extracted and crushed between two frosted glass slides then filtered through a 80 μm filter in FACS buffer. Dissociated spleens were centrifuged at 1200 rpm for 8 minutes at 4°C and the pellet was incubated for 4 minutes in ACK lysis buffer. After the incubation, additional FACS buffer was added and were again centrifuged. Spinal cords were extracted and pushed through 100 μm filters and suspended in a percoll gradient (90%, 37%, and 30%) and centrifuged at 1800 rpm for 20 mins at 4°C. Immune cells were collected at the boundary between the 90% and 37% percoll interface.

If a fixable live/dead was used, cells were washed in PBS once and then stained with Live/Dead Fixable Blue Dead Cell Stain for 30 min on ice. Cells were then washed with FACS buffer and blocked with an anti-Fc-γ receptor for 30 min on ice. Cells were then stained with the primary antibodies targeting surface receptors. Depending on the experiment, cells were either analyzed immediately with the addition of a live/dead green (3 µl in 50 ml of FACS buffer) or cells were fixed. Cells were fixed either by overnight fixation using the Foxp3 / Transcription Factor Staining Buffer Set, fixed for 30 minutes at room temperature using the Intracellular Fixation & permeabilization Buffer set, or fixed for 20 minutes at 4°C in 4% PFA with 1x eBioscience™ Permeabilization Buffer. Samples were blocked with 4% rat serum in 1x eBioscience™ Permeabilization Buffer for 20 min at room temperature, then stained with primary antibodies for intracellular staining. A defined number of CountBright™ Plus Absolute Counting Beads was added to the samples then washed 3 times with 1x permeabilization buffer. Cells were analyzed using a 5-laser Cytec Aurora. Spectral unmixing was done in Spectroflo.

Data was analyzed in FlowJo and for Figures 5G, Supplemental figure 5F-5J, and Supplemental figure 7D-7J the final portions of the analysis were conducted on gated microglia or macrophage in R. Fluorescence values were transformed using an inverse hyperbolic sine (arcsinh) and processed using FlowSOM using the following markers depending on the experiment and cell type: CD68, iNOS, Arg1, CD45, MHC2, CD74, CD115, MerTK, CD11b, CD80, CD64, CD86, CD11c, Ly6C, Tmem119, CD36, CD40, CX3CR1-YFP, PD-L1, SIRPα, and CD206. PCA analysis was then run on the data, followed by harmony for Supplemental figure 7D-7J where multiple experiments were combined, then k-means clustering was used to generate clusters. Clusters were then visualized on a UMAP based on the PCA scores (or harmony correct PCA). Cell abundances and median fluorescence intensity of markers per cluster were then exported from R for analysis in excel.

### Bone marrow derived macrophage generation

Mouse bone marrow derived macrophages were generated as described previously^91^. Briefly, the femurs and tibia were harvested from adult mice, cleaned with gauze and dipped into 70% ethanol for 3 seconds then transferred to DMEM. The edges of the bones were cut and flushed with DMEM using a 25-guage needle. Bone marrow was centrifuged at 500xg for 8 minutes, suspended in PBS and counted. Cells were centrifuged again and suspended at 1.5 x 10^6^ cells per ml in L929 media (10% FBS, 10% supernatant from L929 cells, 1x penicillin and streptomycin, 1x sodium pyruvate, and 1x glutamax), seeded into 35 mm dishes and incubated at 37°C 8.5% CO_2_ for 8 days. Cells were scraped off the bottom of the dishes in ice cold PBS prior to usage.

### Microglia isolation

Mouse microglia were isolated from neonatal brains as described previously^92^ achieving over 95% cell purity. Briefly, the cortices of P0-P2 mouse pups with their meninges removed were diced with a scalpel then vortexed for 1 minute. The cell suspension was pushed through an 80 µm pore filter, then a 11 µm pore filter. Cells were suspended in microglia media (10% FBS DMEM (high glucose) with 1x glutamax, 1x penicillin and streptomycin, 1x sodium pyruvate, and 1x MEM non-essential amino acids) then seeded into T75-flasks such that 1 pup seeds one flask. Cells were cultured at 37°C 5% CO_2_ for 8-10 days then flasks were shaken at 250 rpm for 20 mins in a 37°C orbital shaker to dislodge microglia.

### Transwell assays

700-800 µL of B cell media with BAFF protein (50 ng/ml) or B cell conditioned media was added to the wells of a 24-well plate. In experiments using blocking antibodies, antibodies/isotypes (at a concentration of 5 µg/ml) were added to 50% diluted conditioned media (diluted with B cell media) and incubated at 37°C 5% CO_2_ for 30 minutes. Transwell inserts with 5 µm pores were added to the wells and 200 µL of BMDMs (at a concentration of 1×10^6^ cells per mL in DMEM) was added to the top side of the transwell membrane. Transwells were incubated for 14 or 7 hours (7 hours for blocking experiments) at 37°C 5% CO_2_. Transwells were then removed from their wells, washed in 1 mL of PBS for 5 mins at room temperature, and fixed in acid alcohol (95% ethanol, 5% glacial acetic acid) for 15 minutes at room temperature. Transwells were incubated in hematoxylin for 15 mins at room temperature, washed, and a q-tip was used to remove BMDMs from the topside of the transwell insert. After cleaning, the membranes were cut out of the inserts and mounted in fluoromount-G on glass slides for analysis.

Slides were imaged on a Leica DMI3000B microscope and a Leica DFC320 camera using a 20x lens using brightfield. Two to four fields of view (FOV) across the insert were captured at 25x magnification. The blinded images were analyzed in FIJI using the cell counter module to quantify the number of BMDMs per FOV. The number of BMDMs per field of view was averaged across all images of the same insert.

### In vitro macrophage and microglia stimulation

BMDMs or microglia were seeded into 24-well plates (200000 per well) or 96-well plates (25000 per well) in L929 media or microglia media and incubated at 37°C 5% CO_2_ for 1 day. The next day, media was removed from the wells and washed twice with PBS. 700-800 µL (for 24-well plates) or 90 µL (for 96-well plates) of B cell media with BAFF protein (50 ng/ml) or conditioned B cell media was added to the wells and incubated at 37°C 5% CO_2_ for 1 day. In experiments using blocking antibodies, antibodies/isotypes (at a concentration of 5 µg/ml) were added to 50% diluted conditioned media (diluted with B cell media) and incubated at 37°C 5% CO_2_ for 30 minutes prior to adding them to the wells. After 1 day, the media from each well was removed and the cells were washed with DMEM three times. Then 800 µL (for 24-well plates) or 100 µL (for 96-well plates) of B cell media was added to each well and incubated for 1 day at 37°C 5% CO_2_. Supernatants from the wells were collected, spun at 500 x g for 10 minutes, then frozen at-20°C until needed. For 96-well plates, 100 µL of 4% PFA was added to each well and incubated at room temperature for 20 mins. These cells were stored at 4°C until use. For 24-well plates, 1 mL of FACS buffer with 1mM EDTA was added to the wells and the BMDMs and microglia were scraped off the bottom of the wells. These cells were centrifuged at 500 x g for 8 mins then incorporated into flow cytometry analysis.

### Bulk RNA sequencing and analysis

RNA was isolated from cultured B cells by centrifuging the cells through QIAshredders and isolating RNA using a RNeasy Mini Kit according to the manufacturer’s instructions. Isolated RNA quality and quantity was determined by TapeStation and Fragment Analyzer. Total RNA from each sample was prepared into libraries using the NEBNext Ultra II Directional RNA Library Prep Kit with poly-A enrichment and sequenced using a NextSeq 500 system using a single NextSeq 75-cycle high-output run.

Single-end bulk RNA-seq reads were adapter-trimmed and quality-filtered with Trim Galore and read quality assessed with FastQC. Transcript abundances were quantified by pseudoalignment with kallisto against the Ensembl mouse cDNA reference GRCm39.108 with 100 bootstrap replicates and fragment-length mean/SD of 200/30 for single-end quantification. Gene-level differential expression was performed in sleuth with gene-mode count aggregation using the Wald test and a Benjamini-Hochberg FDR threshold of q < 0.01 and log2 fold change of 1.5, as well as a likelihood ratio test with a FDR of q < 0.05. All analyses were also validated by analyzing the kallisto data using DEseq2 by importing it using tximport.

### Statistics

GraphPad PRISM software was used for statistical comparisons. Non-parametric tests were used for *in vivo* comparisons. Unless otherwise stated, a Mann-Whitney U test was used for comparisons between two groups, and a Kruskal-Wallis test followed by a Dunn’s multiple comparisons post-hoc test was used for comparisons between two groups. For *in vitro* experiments parametric tests were used. A students T-test was used for single comparisons and a one-way ANOVA followed by a Tukey’s multiple comparisons post-hoc test was used for comparisons between more than two groups. Statistical significance is denoted as p*<0.05, p**<0.01, p***<0.001, p****<0.0001.

## Supplemental information

**Supplemental table 1.** Non-neurological control and MS patient characteristics.

**Supplemental table 2.** Differential RNA expression between *in vitro* polarized B cell populations. All differentially expressed genes between polarized B cell cultures (n=4 per group).

---

**Supplemental figure 1.**
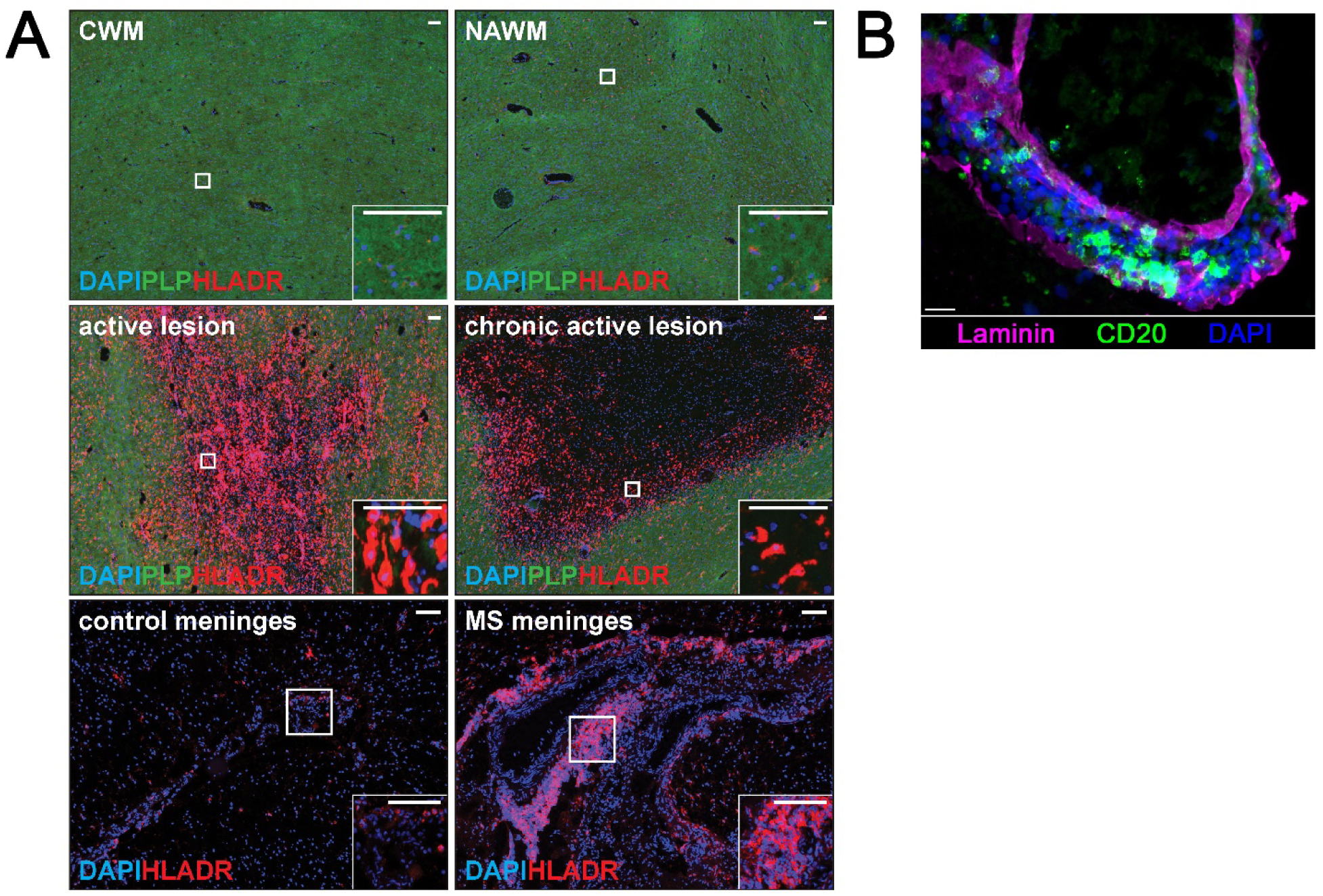
Lesion characterization and additional analyses, related to Figure 1. (A) Representative images of control white matter (CWM), normal appearing white matter (NAWM), active lesions, chronic active lesions, control meninges, and MS meninges showing DAPI (blue), proteolipid protein (PLP) (green), and HLA-DR (red) staining. Scale bars are 100 μm. (B) Representative image of laminin staining outlining the basement membranes of a blood vessel with B cells inside the perivascular space. The lumen of the vessel is the center hole. Scale bar is 20 μm.

**Supplemental figure 2.**
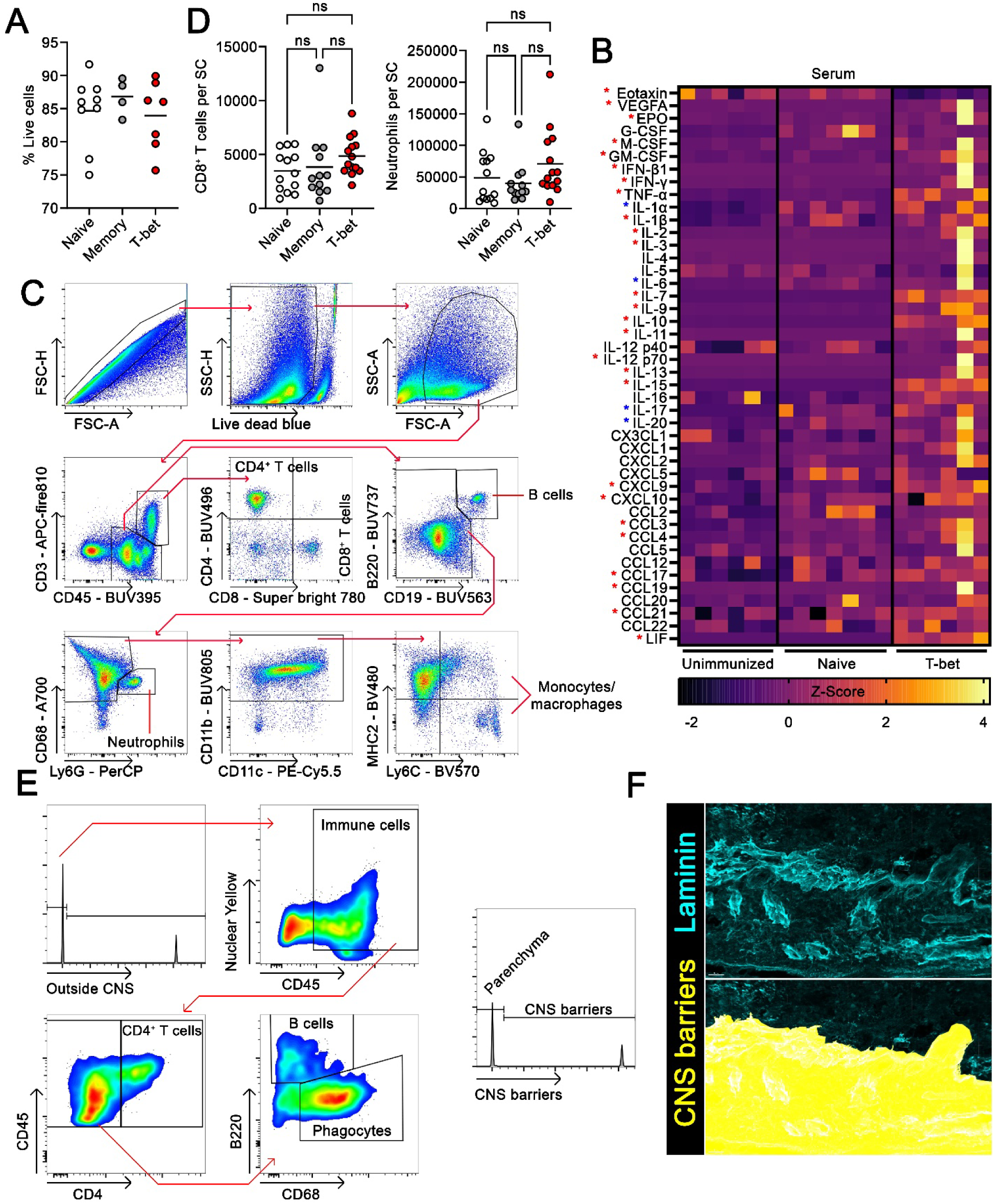
Additional quantification and gating strategies of histoflow and flow cytometry and serum analyses, related to Figure 2. (A) Viability of *in vitro* B cell cultures from various experiments determined by trypan blue staining (n = 4-9). (B) Luminex multiplex analysis of the serum of unimmunized mice or the endpoint of EAE mice with transfer of naïve B cell or T-bet^+^ memory B cells (each box is one Z-scored measurement from one animal; n = 6-7). Blue significance indicates differences between both EAE groups and unimmunized animals and the red significance indicates differences between T-bet and unimmunized only. (C) Gating strategy used for flow cytometry analyses. (D) Quantification of the absolute numbers of CD8^+^ T cells and neutrophils in the spinal cord of the experiment shown in figure 2C (n = 12-14). (E and F) Gating strategy used to identify immune cell populations for histoflow cytometry and an example of how CNS barriers were delineated. Scale bars are 20 μm. Statistical significance was assessed using a one-way ANOVA using Kruskal-Wallis test and Dunn’s post-hoc test (A and D). Data are presented as mean (A and D) or individual values in a heatmap (B) and p-values are summarized as *p<0.05.

**Supplemental figure 3.**
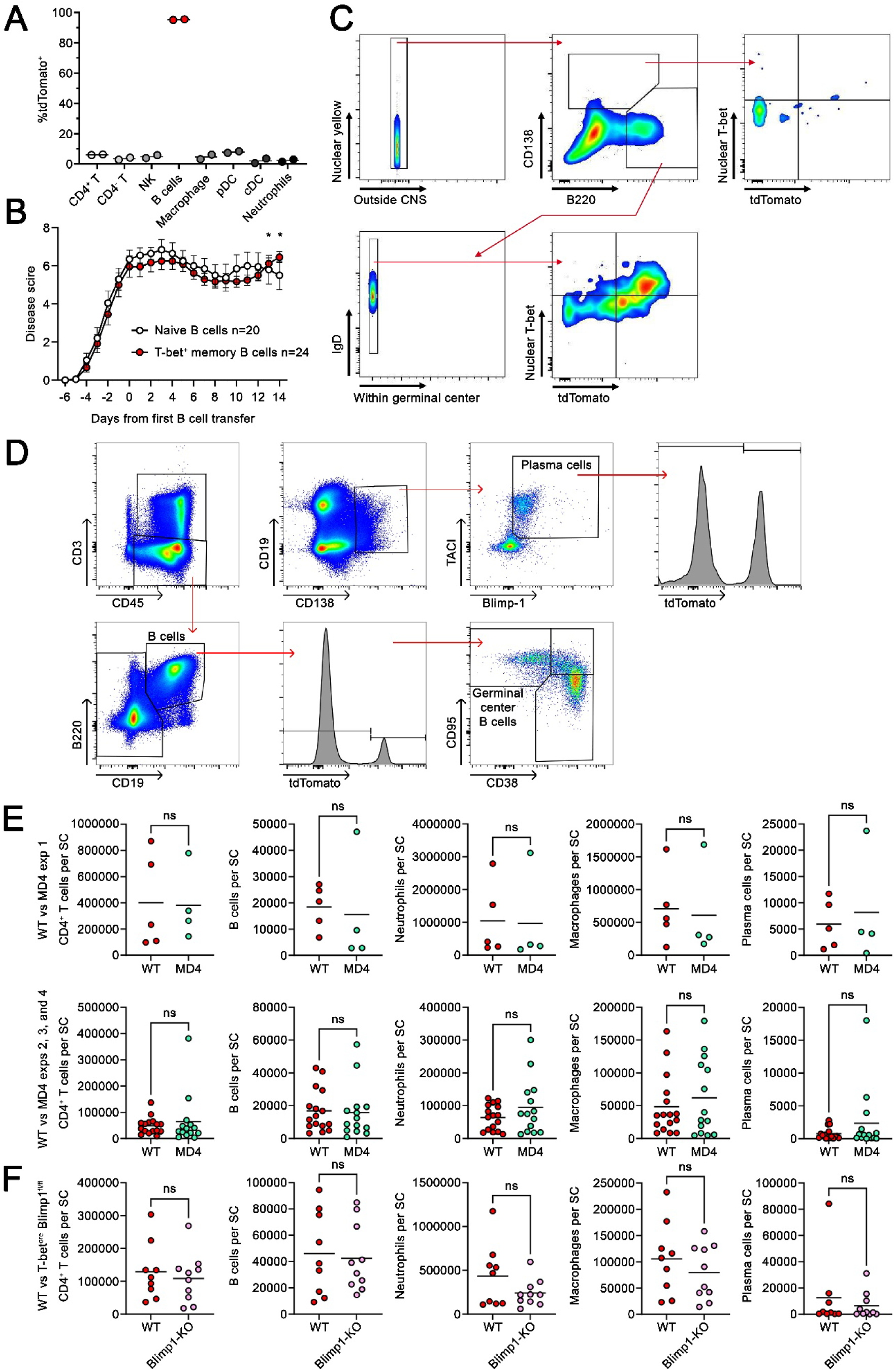
Flow cytometry and histoflow cytometry gating strategies and quantifications of immune cells in the spinal cords of EAE mice, related to Figure 3. (A) Flow cytometry analysis of the blood from Mb1^cre^ x Ai9 mice showing the percentage of the identified populations that are tdTomato^+^. (B) EAE disease score curve comparing EAE mice receiving *in vitro* polarized naïve or T-bet^+^ memory B cell transfers that were differentiated from Mb1^cre^ x Ai9 mice. Combination of 3 independent experiments (n = 20-24). (C) Histoflow cytometry gating strategy used in analysis. (D) Flow cytometry gating strategy used in analysis. (E) Quantification of the absolute number of immune cells entering the spinal cords of EAE mice receiving T-bet^+^ memory B cell transfers derived from wildtype or MD4 mice (n = 4-16). (F) Quantification of the absolute number of immune cells entering the spinal cords of EAE mice receiving T-bet^+^ memory B cell transfers derived from wild type or Blimp1-KO mice (n = 9-10). Statistical significance was assessed using single or multiple Mann-Whitney U tests (B, E, and F). Data are presented as mean ± SEM (B) or mean (A, E, and F) and p-values are summarized as *p<0.05.

**Supplemental figure 4.**
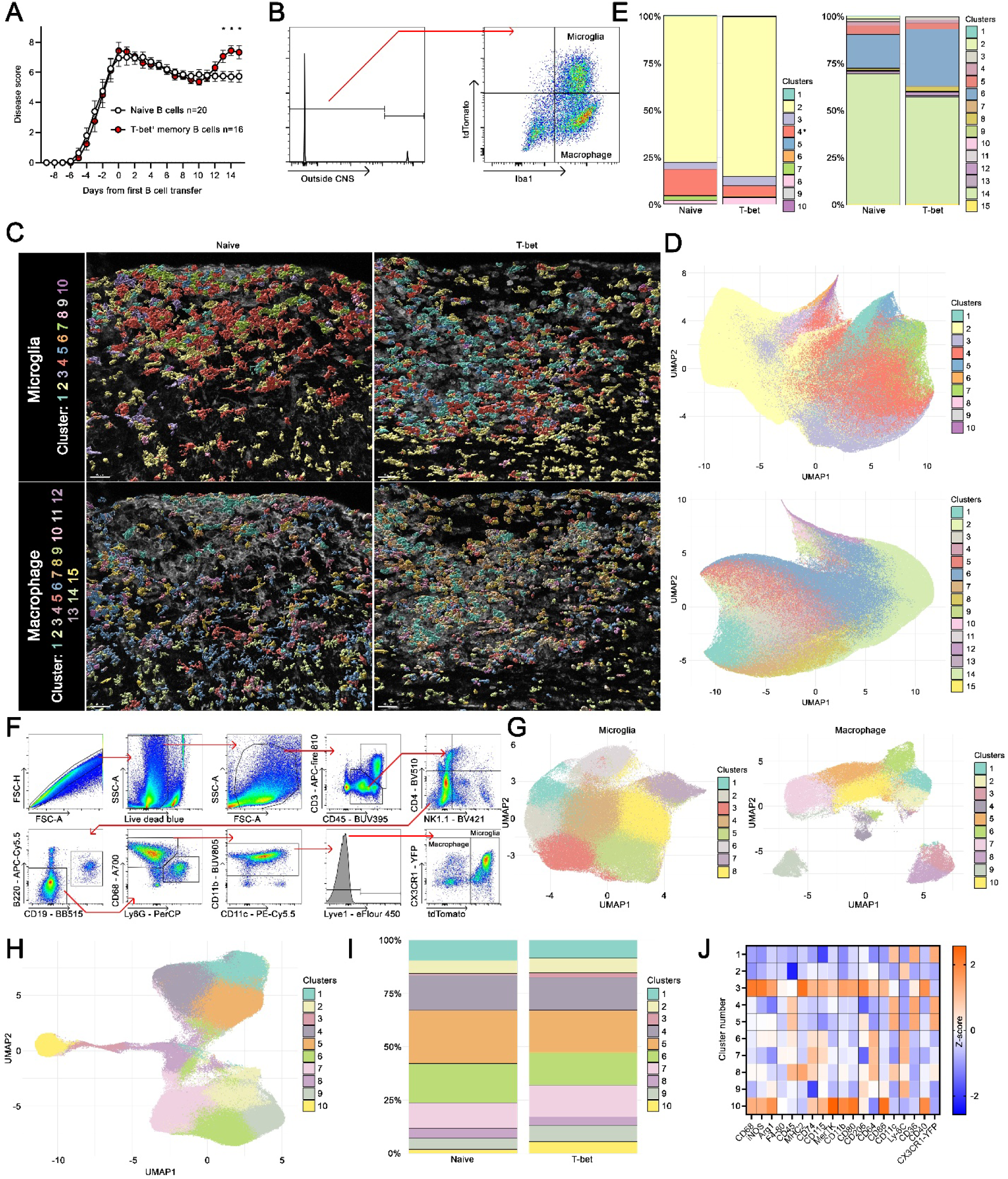
Flow cytometry and histoflow cytometry gating strategies to obtain distinct microglia and macrophage populations and analysis of blood monocytes during EAE, related to Figure 4. (A) EAE disease score curve comparing CX_3_CR1^CreER-YFP^ x Ai9 EAE mice receiving *in vitro* polarized naïve or T-bet^+^ memory B cell transfers. Combination of 3 independent experiments (n = 16-20). (B) The histoflow cytometry gating strategy used to extract cells within the spinal cords separating them into Iba1^+^ tdTomato^-^macrophages and Iba1^+^ tdTomato^+^ microglia. (C and D) UMAPs showing microglia populations (top) and macrophage populations (bottom) are displayed and mapped onto EAE lesions associated with naïve or T-bet^+^ memory B cell transfer. (E) The proportions of microglia (left) and macrophage (right) populations in white matter outside of lesions is shown for EAE mice receiving naïve or T-bet^+^ memory B cell transfer (n = 4-5). (F) Representative gating strategy used to isolate macrophage and microglia in the spinal cords of CX_3_CR1^CreER-YFP^ x Ai9 EAE mice receiving *in vitro* polarized naïve or T-bet^+^ memory B cell transfers. (G) UMAPs showing microglia (left) and macrophage populations (right) in the CNS. (H) UMAP showing macrophage populations in the blood of CX_3_CR1^CreER-YFP^ x Ai9 EAE mice. (I) Proportions of macrophage populations in the blood of EAE mice receiving naïve or T-bet^+^ memory B cells and a heatmap showing Z-scored expression of the indicated molecules (J) (n = 6-8). Representative 1 of 2 experiments. Statistical significance was assessed using single or multiple Mann-Whitney U tests (A, E, and I). Data are presented as mean ± SEM (A) or are only shown as proportion (E and I) and p-values are summarized as *p<0.05.

**Supplemental figure 5.**
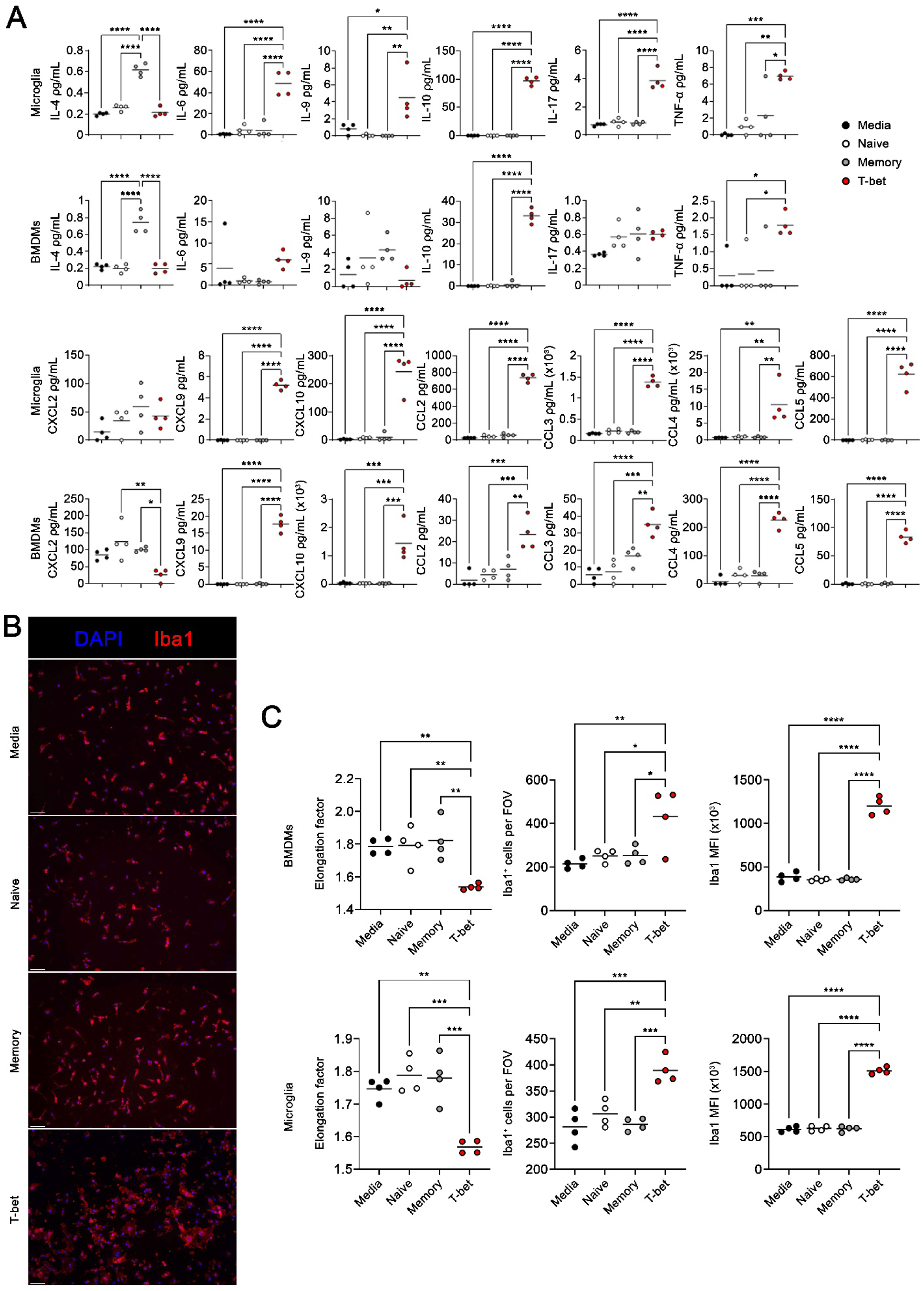
Quantification of cytokine and chemokine expression from stimulated BMDMs and microglia and associated morphological analysis, related to Figure 5. (A) Quantification of Luminex analyses from Figure 5F in pg/ml (n = 4). Representative 1 of 2 experiments. (B and C) Imaging and analysis of BMDMs and microglia after stimulation with B cell supernatants and incubation in blank media for 24 hrs. (B) Representative images of microglia stimulated with B cell supernatants. (C) Quantification of how elongated (left), cell numbers (middle), and Iba1 MFI of BMDMs (top) and microglia (bottom) after stimulation with B cell supernatants (n = 4). Representative of one experiment. Statistical significance was assessed using a parametric one-way ANOVA with a tukey’s post-hoc test (A and C). Data are presented as mean (A and C) and p-values are summarized as *p<0.05, **p<0.01, ***p<0.001, and ****p<0.0001.

**Supplemental figure 6.**
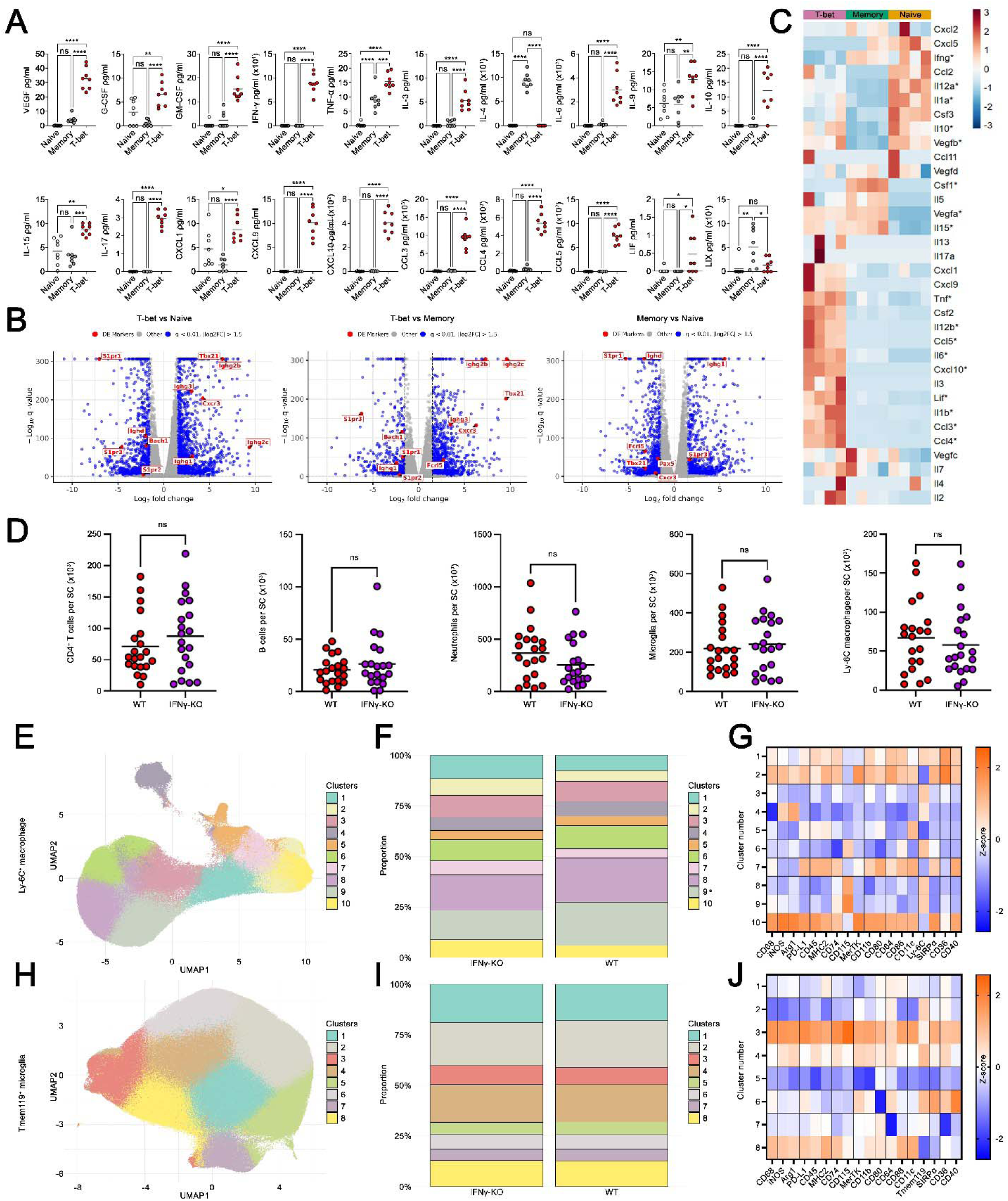
Bulk RNA sequencing analysis and quantification of cytokine and chemokine expression from polarized B cells and analyses of immune cell composition in the CNS of EAE mice receiving wildtype or IFNγ-KO mice, related to Figure 6. (A) Quantification of Luminex analyses from Figure 6A in pg/mL (n = 8). Combination of two independent experiments. (B and C) B cells from wild type animals were *in vitro* polarized into naïve, memory, or T-bet^+^ memory B cells for 4 days then harvested for bulk RNA sequencing. (B) Volcano plots showing differentially expressed genes between populations. (C) A heatmap showing Z-scored transcripts per million of chemokines and cytokines between the B cell subsets (n = 4). Based on one experiment. (D) Quantification of immune cell infiltration into the spinal cords of the mice from Figure 6L by flow cytometry (n = 20-22). Combination of three independent experiments. (E) UMAP showing Ly-6C^+^ Tmem119^-^ macrophage subsets determined by unsupervised clustering. (F) Proportions of macrophage subsets in the spinal cords of EAE mice receiving wildtype or IFN-γ-KO T-bet^+^ memory B cells. Combination of three independent experiments. (G) Heatmap showing the Z-scored expression of the displayed molecules across the macrophage subsets. (H) UMAP showing Ly-6C^-^ Tmem119^+^ microglia subsets determined by unsupervised clustering. (I) Proportions of microglia subsets in the spinal cords of EAE mice receiving wildtype or IFN-γ-KO T-bet^+^ memory B cells. Combination of three independent experiments. (J) Heatmap showing the Z-scored expression of the displayed molecules across the microglia subsets. Statistical significance was assessed using a parametric one-way ANOVA with a tukey’s post-hoc test (A), a Wald test and a Benjamini-Hochberg FDR threshold of q < 0.01 and log2FC of 1.5 (B), a likelihood ratio test with a FDR of q < 0.05 (C), or a Mann-Whitney U test (D, F, and I). Data are presented as mean (A and D), individual values (C), or are only shown as proportion (F and I) and p-values are summarized as *p<0.05, **p<0.01, ***p<0.001, and ****p<0.0001.

**Supplemental figure 7.**
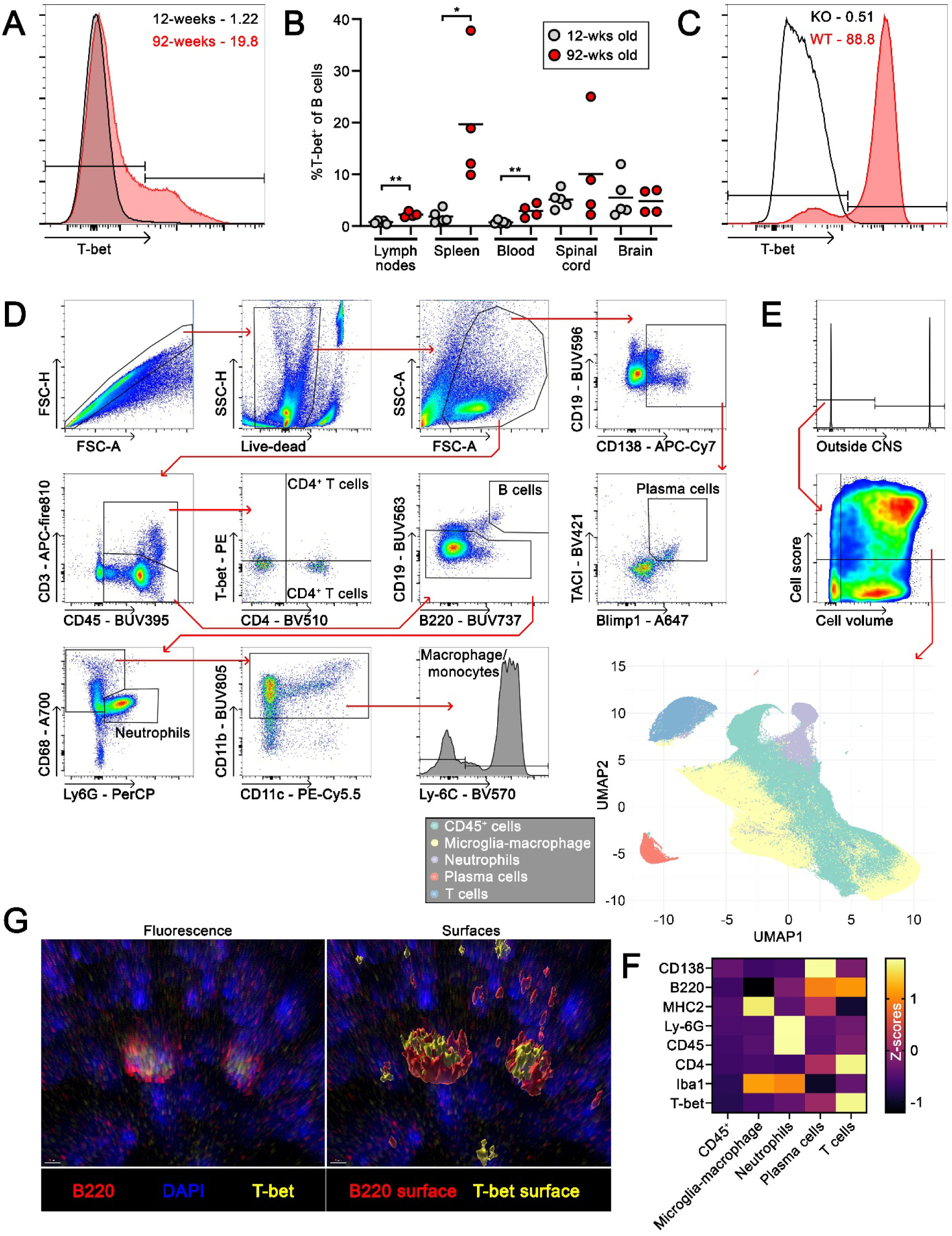
Flow cytometry and histoflow cytometry gating strategies for evaluating immune cell compositions in the CNS of aged or young T-bet wild type or knockouts and knockout efficiency, related to Figure 7. (A and B) T-bet expression in CD19^+^ B220^+^ B cells in the spleens of 12- or 92-week old mice shown in a histogram (A) and quantified in (B) (n = 4-5). (C) A histogram showing T-bet expression in *in vitro* polarized B cells (polarized into T-bet^+^ memory B cells) from Mb1-cre^+/-^ Tbx21^fl/fl^ mice (KO) or Mb1-cre^-/-^ Tbx21^fl/fl^ mice (WT). (D) Representative gating strategy used to identify plasma cells, B cells, CD4^+^ T cells, neutrophils, and macrophage/monocytes in the spinal cords WT or KO EAE mice that were young or old. (E) Histoflow cytometry gating strategy used to extract quality cells in the spinal cords before conducting quality control and unsupervised clustering to yield the UMAP shown below identifying 5 major populations. (F) A heatmap showing the Z-scored expression of the indicated markers in each population from E. (G) Representative images from the spinal cord of a WT old EAE mouse showing T-bet staining within B220 staining (left) and surfaces created over these stains to show the localization of T-bet wrapped within B220. Scale bars are 5 μm. Statistical significance was assessed using a student’s T-test (B). Data is presented as mean (B) and p-values are summarized as *p<0.05, and **p<0.01.

## Notes

### Competing Interest Statement

The authors have declared no competing interest.

